# Host-Specific Adaptations and Global Transmission Potential of a Mexican ToBRFV Isolate: Insights from Genomic Analysis

**DOI:** 10.1101/2025.01.08.631997

**Authors:** Zamora-Macorra Erika Janet, Ochoa-Martínez Daniel Leobardo, Chavarín-Camacho Claudia Yaritza, Rosemarie W Hammond, Aviña-Padilla Katia

## Abstract

The tomato brown rugose fruit virus (ToBRFV) poses a severe global threat to tomato and pepper production due to its high transmissibility and genetic adaptability. This study provides an in-depth characterization of a Mexican ToBRFV isolate, integrating genomic variability analysis, host-specific investigations, and seed transmission studies to better understand the virus’s adaptability and dissemination pathways. Phylogenetic analysis places this Mexican isolate in close genetic proximity to strains from Israel and China, suggesting cross-regional transmission potentially facilitated by agricultural trade. Genomic comparison with 100 ToBRFV isolates revealed a notable balance between genomic stability and host-specific adaptations, particularly within the 126-kDa replicase domain proteins critical for viral replication. Specific single nucleotide variants (SNVs) were identified in these regions, exhibiting host-dependent patterns; *Solanum lycopersicum* (tomato) isolates maintained consistent profiles, whereas isolates from *Capsicum annuum* (pepper) and *Citrullus lanatus* (watermelon) displayed unique substitutions. These variations suggest distinct selection pressures across hosts, potentially optimizing viral replication and survival in specific plant environments. Experimental bioassays confirmed that the Mexican isolate could infect additional species, including *Physalis ixocarpa*, and *Solanum melongena*, thereby expanding the known host range of ToBRFV. Seed transmission studies conducted on *N. rustica* revealed a 30% reduction in germination rates for infected seeds compared to healthy controls, highlighting the virus’s impact on seed viability and underscoring the necessity of stringent biosecurity protocols. This comprehensive analysis enhances our understanding of the Mexican ToBRFV isolate genetic diversity, host specificity, and transmission dynamics.

## 1 Introduction

Tomato Brown Rugose Fruit Virus (ToBRFV), an emergent tobamovirus, poses a critical threat to global agriculture, particularly affecting essential Solanaceae crops like tomato (*Solanum lycopersicum*) and pepper (*Capsicum annum*) (Fidan et al., 2020; Magaña-Álvarez et al., 2021). This rod-shaped, positive-sense single-stranded RNA virus (+ssRNA) is notable for its robustness and high transmissibility, with rapid spread reported across key agricultural regions (Luria at el., 2017). In greenhouse settings, ToBRFV transmission is predominantly mechanical, facilitated by routine agricultural practices that inadvertently promote viral spread (Panno et al., 2020).

The genome of ToBRFV consists of approximately 6,400 nucleotides organized into four essential open reading frames (ORFs) (Salem *et al*., 2023). This genomic architecture is central to ToBRFV’s infectivity, adaptability, and spread among plant hosts, particularly economically significant crops. The viral replication machinery is formed by the proteins encoded in ORF1 and ORF2, namely the 126 kDa and 183 kDa replicase-associated proteins. ORF1 directly encodes the 126-kDa protein (p126), which includes methyltransferase and helicase domains necessary for viral RNA synthesis and host immune suppression, as noted in related tobamoviruses (Ishibashi and Ishikawa, 2016.). A readthrough mechanism of the stop codon between ORF1 and ORF2 produces the 183-kDa protein (p183), containing a C-terminal RNA-dependent RNA polymerase (RdRp) domain essential for viral replication. ToBRFV’s systemic infection capacity is driven by ORF3 and ORF4, which are translated from subgenomic RNAs (sgRNA1 and sgRNA2, respectively). ORF3 encodes a 30-kDa movement protein (MP) that enables cell-to-cell viral spread by modifying plasmodesmata, ensuring effective systemic infection. ORF4 encodes the 17.5-kDa coat protein (CP), which encapsulates the viral RNA, thereby enhancing stability, protecting against degradation, and facilitating both intracellular and inter-host transmission (Salem et al., 2016). The 5′ untranslated region (UTR), rich in CAA repeats, and the 3′ UTR with tRNA-like structures further support ToBRFV’s resilience and adaptability across different host plants.

Genomic and evolutionary analyses are essential for understanding the transmission pathways and host adaptations of the virus. These studies enable researchers to identify adaptive mutations, trace transmission routes, and infer evolutionary relationships that facilitate the virus’s establishment across diverse hosts and environmental conditions (Rizzo et al., 2021; Abrahamian et al., 2022; Zisi et al., 2024). Such information is critical in designing targeted containment strategies tailored to the unique genetic characteristics of local ToBRFV strains.

Recent genomic studies reveal that ToBRFV exhibits remarkably low variability, supporting a monophyletic origin, with over 99% nucleotide identity shared across all known genomic sequences (Abrahamian et al., 2022; Çelik et al., 2022). Limited single-nucleotide polymorphisms (SNPs) have been identified, showing a relatively even distribution across the genome without distinct hotspots of variation (Van de Vossenberg et al., 2021). This high sequence conservation and limited variability underscore ToBRFV’s rapid global spread from a single origin, likely driven by international trade in tomato seeds and fruits prior to the implementation of stringent control measures. A recent observation in the Netherlands highlights the emergence of a novel phylogenetic clade, potentially resulting from the unauthorized use of an isolate for cross-protection. This finding underscores the importance of strict adherence to quarantine regulations to effectively manage this pathogen and prevent further spread (Salem et al., 2023; Botermans et al., 2023).

ToBRFV has rapidly become a focus of agricultural biosecurity efforts worldwide due to its resilience across diverse environments (<u>Datab</u>https://gd.eppo.int/taxon/TORFV/distributions). Mexico, with its rich biodiversity in Solanaceae species, including endemic varieties, has emerged as a hotspot for ToBRFV transmission (Vargas-Mejía et al., 2023; García-Estrada et al.,2022). The country’s unique agricultural and ecological dynamics make it a critical region for studying ToBRFV’s adaptation and transmission. Characterizing local isolates is essential to understanding how region-specific adaptations may arise in response to local agricultural practices, crop varieties, and trade dynamics.

Since its detection in Mexico in 2018 (Rodríguez-Mendoza et al., 2019), ToBRFV has established itself across all major tomato-growing regions, raising concerns about its ability to overcome resistance genes (*Tm-1, Tm-2,* and *Tm-22*) traditionally used in tomato cultivars, leading to significant crop losses in high-density greenhouse systems. A recent second introduction of ToBRFV into Mexico, related to strains from the Netherlands and the Middle East, underscores the virus’s global transmission risk, likely facilitated by international trade. Seed coat and epicotyl RT-PCR testing revealed a 9% transmission rate, highlighting the role of seeds in viral dissemination (Vargas-Mejía et al., 2023).

Although tomato and pepper are ToBRFV’s primary natural hosts, experimental inoculations reveal the virus’s ability to infect a broad range of species, including *Chenopodium* and *Nicotiana* spp., demonstrating remarkable adaptability. A key factor in this adaptability is ToBRFV’s genomic plasticity, which allows host-specific mutations that enhance viral fitness in economically significant hosts such as tomato and pepper, as well as in endemic species like *S. nigrum* (Salem et al., 2022), *S. eleagnifolium*, *S. rostratum* (Matzrafi et al., 2023), *Ipomoea purpurea, Mirabilis jalapa,* and *Clematis drummondii* (Vasquez-Gutierrez et al., 2024). These host-driven mutations, shaped by selective pressures, improve ToBRFV’s infectivity and replication efficiency, allowing it to persist across cultivated and wild plant reservoirs. This flexibility underscores the need for targeted management strategies that address ToBRFV’s impact in both agricultural and natural settings, particularly in biodiverse regions such as Mexico, where endemic species may act as reservoirs.

ToBRFV spreads mechanically and through seeds, where it primarily resides on the seed coat and endosperm, without affecting the embryo. Although seed-to-seedling transmission rates are relatively low (0.08% to 1.8%), the virus remains a biosecurity concern, especially when micro-wounds in seedlings facilitate infection (Salem et al., 2022; Davino et al., 2020). Seed-borne transmission is particularly challenging as it allows the virus to bypass containment protocols and spread across borders, threatening global trade and introducing ToBRFV to new regions. This transmission route emphasizes the need for rigorous seed surveillance and treatment protocols to contain ToBRFV and safeguard agricultural productivity.

Herein we provide a detailed characterization of a Mexican ToBRFV isolate, emphasizing host-specific mutations, adaptive genetic traits, and seed-borne transmission potential. By integrating genomic analysis, experimental validation, and biosecurity considerations, this research advances understanding of ToBRFV’s adaptability and transmission dynamics, addressing both regional and global challenges posed by this significant agricultural pathogen.

## 2 Materials and Methods

### 2.1 Location and Field Sampling

In September 2020, tomato plants *(Solanum lycopersicum* L.) exhibiting classic symptoms of ToBRFV infection - including chlorosis, leaf narrowing, and mosaics - were sourced from commercial greenhouses in Colima Mexico. Subsequent RT-PCR testing confirmed the presence of ToBRFV in the collected samples.

### 2.2 Genome sequencing and Analysis workflow

RNA extraction was performed using CTAB2%-Trizol® protocol following the protocol described by Jordon-Thaden et al. (2015) (minor modifications), followed by library preparation involving fragmentation and adapter ligation. The libraries were amplified, purified, and sequenced using Illumina’s Sequencing by Synthesis (SBS) technology by Psomagen®. Raw sequencing data in BCL format was converted to `FASTQ` files for downstream analysis. Quality control on the raw reads included assessments of total bases and GC content, along with trimming to remove adapter sequences. The cleaned reads were assembled *de novo* using `SPADES`, with parameters optimized for contig number and N50. Assembly accuracy was validated by mapping the reads back to the genome, ensuring precise genomic reconstruction. Functional annotation was performed using the conserved domains tool (https://www.ncbi.nlm.nih.gov/Structure/cdd/wrpsb.cgi) (Wang et al., 2023; Lu et al., 2020; Marchler-Bauer et al., 2017). Then, we implemented a comprehensive bioinformatics workflow to analyze the genome of the Mexican ToBRFV isolate. The workflow is illustrated in **Supplemental Figure 1**, and all code used in these analyses is accessible in a GitHub repository at https://github.com/kap8416/TOBRFV-Genome-Analyses/tree/main. The full genome sequences of 100 ToBRFV isolates were retrieved from the GenBank database (https://www.ncbi.nlm.nih.gov/genbank/, accessed in July 2024). Metadata, including geographical location and host information, was systematically extracted and organized using custom scripts in the `RStudio` environment (https://www.rstudio.com/). This comprehensive dataset provided a foundation for investigating the virus genome characteristics and distribution patterns.

### 2.3 Phylogenetic Analysis

Initially, 101 ToBRFV genome sequences were aligned using the `ClustalW` algorithm, with the input sequences in `FASTA` format. This alignment was then used as input for phylogenetic analysis in `IQ-TREE2` using bash scripts. The analysis employed the `Maximum Likelihood` (ML) method, with `ModelFinder` automatically selecting the best-fitting substitution model. To evaluate clade support, 1000 ultrafast bootstrap replicates were applied. The resulting phylogenetic tree was visualized in `Rstudio` using the `ggtree` package for enhanced visualization. A pairwise sequence identity matrix was generated using the `ape` package to quantify nucleotide identity percentages between isolates, and visualized as a heatmap with the `pheatmap` package. Then, a Neighbor-Net phylogenetic network was generated using the `phangorn` and `ape` packages to examine relationships among the ToBRFV isolates. Finally, key genetic diversity parameters, including haplotype diversity (Hd), the number of haplotypes (H), nucleotide diversity (π), and the average nucleotide differences (k), were calculated with the `pegas` and `ape` packages. A radar plot was created with the `fmsb` package to visually compare these diversity indices across isolates.

### 2.4 Genomic Variation and Distribution Analysis of the Mexican ToBRFV Isolate

#### 2.4.1 Nucleotide Variation Analysis

To detect and visualize specific genomic differences between the Mexican ToBRFV isolate and 100 reference genomes, a nucleotide variation analysis was performed using the `Biostrings` package in RStudio. Alignment with reference genomes enabled identification of nucleotide positions where the Mexican isolate diverged. To ensure consistent variability measurement, observed differences at each position were normalized for sequence length and alignment quality. Statistical summaries, including mean and standard deviation, were computed to highlight regions with higher or lower mutation rates. Visualization of normalized difference frequencies across the genome was achieved using `ggplot2,` with elevated values pinpointing sites of increased variability.To analyze sequence variability and complexity in the Mexican ToBRFV isolate, Shannon entropy was calculated for each nucleotide position based on alignment data, **Supplemental Figure 4**. Entropy values were plotted alongside genomic differences to identify regions of high diversity and potential functional significance. Sliding window analyses smoothed trends, and correlation metrics quantified the relationship between entropy and mutation frequency. These approaches provided insights into sequence diversity and its genomic distribution. Correlation metrics, including Pearson and Spearman coefficients, quantified the association between mutation frequency and entropy. Data was analyzed and visualized using custom scripts in `RStudio.`

#### 2.4.2 Geographical Distribution of ToBRFV Isolates

Geographical metadata associated with each ToBRFV isolate was extracted and visualized using `ggplot2` (version 3.3.2) and `sf` (version 0.9-6) packages in `RStudio`. A world map annotated with counts of genetically similar isolates from each region provided a visual overview of the global distribution and regional clustering of ToBRFV isolates closely related to the Mexican genome, illustrating the global dissemination patterns of the virus.

#### 2.4.3 Alignment Scores by Host and Geographical Distribution

To examine alignment scores in relation to natural hosts and geographical distribution, alignment scores for each sequence were calculated using the `msa` package (version 1.22.0). These scores were combined with host and geographic data to create an integrated dataset. A scatter plot generated with `ggplot2` displayed alignment scores, with points colored by host and annotated by region, illustrating alignment score variations across hosts and geographical locations.

#### 2.4.4 Host-Specific Nucleotide and Amino Acid Differences

To explore nucleotide variation associated with different hosts, sequence variants were identified across host groups using the `Biostrings` package. These differences were integrated with host data using creating a dataset linking genetic variation to host information. Visualization with `ggplot2` revealed patterns of nucleotide differences along the genome, highlighting potential host- specific adaptations and evolutionary pressures exerted by different plant hosts on ToBRFV.The nucleotide sequence of the ToBRFV Mexican isolate was preprocessed to ensure continuity for analysis. Codon and amino acid translations were performed using a custom R script with the `seqinr` package to map nucleotide positions to codons and amino acids. The methyltransferase (280–1365) and helicase (2543–3315) domains, critical for replication and transcription, were analyzed to evaluate the impact of single nucleotide variants (SNVs). Mutation effects were categorized as *“Change”* or *“No change”* based on amino acid substitutions. Using `ggplot2`, scatterplots visualized SNVs.

### 2.5 Experimental Validation of ToBRFV Hosts

#### 2.5.1 Inoculum Preparation, RNA Extraction and RT-PCR Confirmation

To prepare the inoculum, a *Nicotiana glutinosa* plant was initially infected with ToBRFV-positive tomato tissue, resulting in necrotic lesions. This infection was sequentially transferred through four generations of *N. glutinosa* plants to ensure consistency. The final inoculum was then used to infect tomato plants. This inoculum was used for further host bioassays and transmission studies. Plant tissues were first mechanically macerated in phosphate buffer (0.1 M potassium phosphate, pH 7.0; 0.1 M sodium sulfite; 1% β-mercaptoethanol) and used to inoculate uninfected tomato seedlings which were confirmed to be infected via RT-PCR at 30 days post-inoculation (dpi). Infected plant tissues were processed for RNA extraction and ToBRFV presence was confirmed by RT-PCR, following protocols adapted from Rodríguez-Mendoza et al. (2019). Resulting amplicons were sequenced to verify viral identity.

#### 2.5.2 Host Bioassays

To identify potential ToBRFV hosts, mechanical inoculations were performed on five biological replicates from each of the following species; a) Solanaceae family: tomato *(Solanum lycopersicum L.)*, tomatillo *(Physalis ixocarpa)*, tobacco *(Nicotiana rustica)*, pepper *(Capsicum annum)*, eggplant *(Solanum melongena L.)*; b) Cucurbitaceae family; watermelon *(Citrullus lanatus)*, cantaloupe (*Cucumis melo L.)*, squash (*Cucurbita pepo*), and cucumber (*Cucumis sativus L.*). Additionally, three plants from each species were mock-inoculated as controls. The plants were maintained under greenhouse conditions and observed daily to record symptoms. After 30 dpi, plants were tested for virus presence using RT-PCR employing the primers described by Dovas et al. (2004) and ELISA, with antibodies against TMV (Agdia®) that have cross-reactions. This experiment was performed thrice: first in October 2019, second in February 2020, and last in April 2020. During the first experiment, the average monthly temperature in the greenhouse ranged from 13°C to 23°C with a 12 hours light:12 hours dark cycle. During the second experiment, the temperature went from 11° to 26° with an 11 hours light:13 hours dark cycle. The temperature during the third experiment ranged from 14° to 29°C with a 13 hours light:11 hours dark cycle.

### 2.6 Seed Transmission Test: Acquisition of Seeds

For our seed transmission test, thirty healthy *N. rustica* tobacco plants at the two true leaves stage were subjected to mechanical inoculation with the virus. Simultaneously, a separate group of plants was mock-inoculated to act as controls. Following inoculation, all plants were cultivated under stringent greenhouse conditions, with a stable 12-hour light:12-hour dark photoperiod, and temperatures maintained between 16°C and 24°C. As the plants reached fruiting, seeds were carefully harvested from each once their capsules opened. These seeds were then stored at a cool 4°C for preservation. To validate the successful transmission and presence of ToBRFV in the tobacco plants, a post-harvest assessment was carried out. This involved extracting total RNA from the plants and performing RT-PCR analysis, with the specific primers derived from the methodology presented by Dovas et al. (2004).

#### 2.6.1 Seed Germination Test

Utilizing stereoscopic microscopy, empty seeds were meticulously identified and set aside. These were further segregated into three distinct batches, each containing 100 seeds sourced from ToBRFV-infected plants, alongside seeds from mock-inoculated controls. For germination, each seed batch was placed in a petri dish lined with moistened paper, using sterile distilled water as the wetting agent. The petri dishes underwent an initial incubation at 23°C, exposed to continuous light for 72 hours. This was succeeded by a shift to a 12-hour light: 12-hour dark cycle with temperatures oscillating between 18°C and 24°C. After a span of 20 days, the germinated seedlings in each dish were enumerated. This entire germination regimen was conducted in triplicate to ensure reproducibility.

#### 2.6.2 Estimation of Percentage of Infection in Seeds or Seedlings and experimental method to determine percentage of infected seeds

To gauge the infection rate in seeds or seedlings, we adopted the method propounded by Albrechtsen (2006). This approach hinges on the maximum likelihood estimation. Specifically, our entire sample was segmented into distinct subgroups (N), each with a defined number (n) of specimens, **Supplemental Figure 2.** Employing the formula P=[1−(Y/N)1/n]×100*P*=[1−(*Y*/*N*)1/*n*]×100, where ‘Y’ signifies the number of infection-negative subgroups, we determined the infection percentage. This method was then precisely applied to ascertain the infection rate in *N. rustica* seedlings sprouted from seeds originating from ToBRFV-infected plants, providing a comprehensive view of ToBRFV’s prevalence within our samples.

To assess the percentage of ToBRFV-infected seeds, we initiated with fifteen subsets of 250 seeds each, germinated in Petri dishes under predefined conditions. Using stereoscopic microscopy, sprouted seedlings were isolated, separating the seed coat from the cotyledon. These seedlings, stored at -20°C in sterilized envelopes, totaled 2,250 post-collection. Employing the CTAB-Trizol method from Jordon-Thaden et al. (2015), total RNA was extracted, standardized to 300 ng/µL, and utilized for retrotranscription. Nested PCR, adhering to protocols by Dovas et al. (2004), was executed, followed by 1.5% agarose gel electrophoresis to verify fragment sizes. Purified products underwent sequencing at Macrogen, Korea. A parallel effort involved 15 subsets of 150 *N. rustica* seeds from ToBRFV-positive plants, processed similarly. Both assays incorporated controls—two subsets of 150 seeds or seedlings from healthy *N. rustica*. This comprehensive approach aimed to accurately ascertain the extent of ToBRFV infection in seeds.

#### 2.6.3 Infectivity tests of viral particles in seeds

We performed infectivity evaluations utilizing fifteen subsets, each containing 100 seeds. These were treated with 3% sodium hypochlorite for three minutes, with parallel tests on untreated seed subsets for comparison. Post-treatment, seeds from each group were macerated in liquid nitrogen. This macerate was apportioned equally between two Eppendorf tubes; the first with 500 µL of phosphate buffer (pH 7.0) and the second filled with 600 µL of 2% CTAB buffer. Subsequently, healthy *N. rustica* plants were mechanically inoculated, using carborundum as an abrasive, and then subjected to CTAB RNA extraction and nested RT-PCR (**Supplemental Figure 3**). Inoculated *N. rustica* plants were nurtured in a greenhouse setting, with temperatures ranging between 18°C and 23°C, and maintained under a 12-hour light-dark cycle. Daily observations were made to monitor symptom emergence. By 30 dpi, apical leaves from each specimen were chosen for total RNA extraction and subsequent RT-PCR analysis. Control groups encompassed two subsets of 100 treated seeds, 100 untreated seeds, and 100 seedlings, all sourced from healthy *N. rustica*.

## 3 Results

Tomato plants (*Solanum lycopersicum* L.) exhibiting classic symptoms of ToBRFV infection— chlorosis, leaf narrowing, and mosaic patterns—were collected from commercial greenhouses in Mexico. Reverse transcription PCR (RT-PCR) confirmed the presence of ToBRFV in the symptomatic samples. Sequencing of the Mexican ToBRFV isolate produced 50,136,646 raw reads with a GC content of 50.15% and high-quality scores (Q20: 97.03%, Q30: 92.37%) (**Supplemental material**). After quality trimming, 44,518,584 reads remained, retaining a GC content of 50.35% and achieving improved quality scores (Q20: 98.93%, Q30: 95.61%). Further filtering yielded a final dataset of 99,338 reads, with a GC content of 50.34%.

Mapping analysis revealed that 31.41% of the reads aligned to the reference genome, achieving 100% coverage with an average depth of 607.16×. Contig assembly generated a single contig of 6,375 bases with a GC content of 41.54%, demonstrating high sequencing and assembly fidelity. These results provide a solid foundation for subsequent genomic and computational analyses to elucidate the genetic characteristics and transmission dynamics of this Mexican ToBRFV isolate.

The annotated genome of the Mexican ToBRFV isolate revealed key conserved protein domains essential for viral replication, transcription, and host interaction (**Table 1**). The viral *methyltransferase* domain (280–1365, pfam01660, E-value: 9.92E-08) is critical for RNA capping, enhancing RNA stability and host translation compatibility. The *viral helicase* domain (2543–3315, pfam01443, E-value: 4.31E-19) supports RNA unwinding during replication and transcription.

**Table 1.** Annotated Functional Domains in the Genome of the Mexican ToBRFV Isolate.

| Name | Accession | Description | Interval (nucleotides) | E-value |
| --- | --- | --- | --- | --- |
| **Viral Methyltransferase** | pfam01660 | Viral methyltransferase; This RNA methyltransferase domain is found in a wide range of ssRNA ... | 280-1365 | 9.92E-08 |
| **Viral_helicase1** | pfam01443 | Viral (Superfamily 1) RNA helicase; Helicase activity for this family has been demonstrated ... | 2543-3315 | 4.31E-19 |
| **Virgaviridae_RdRp** | cd23251 | RNA-dependent RNA polymerase (RdRp) in the family Virgaviridae of positive-sense ... | 3514-4803 | 2.02E-97 |
| **RdRP_2** | pfam00978 | RNA dependent RNA polymerase; This family may represent an RNA dependent RNA polymerase. The ... | 3565-4887 | 1.04E-96 |
| **MP** | pfam01107 | Viral movement protein (MP); This family includes a variety of movement proteins (MPs). The MP ... | 4907-5392 | 8.28E-25 |

Two overlapping RNA-dependent RNA polymerase (RdRp) domains were identified (*Virgaviridae_RdRp*, 3514–4803, cd23251, E-value: 2.02E-97; *RdRP_2*, 3565–4887, pfam00978, E-value: 1.04E-96), critical for viral RNA synthesis. The viral movement protein (MP) domain (4907– 5392, pfam01107, E-value: 8.28E-25) enables intercellular virus transport through plasmodesmata. These findings confirm the presence of highly conserved and functionally significant domains essential for the pathogenicity and life cycle of ToBRFV.

### 3.1 High genetic Similarity and Limited Divergence of the Mexican ToBRFV Isolate Among Global Variants

To place the Mexican ToBRFV isolate within a global evolutionary framework, we conducted a comprehensive comparative analysis against 100 ToBRFV isolates. This approach allowed us to understand the isolates genetic relationships, potential transmission routes, and evolutionary stability. Our results revealed that the Mexican isolate closely clusters with other isolates, displaying high genetic similarity and suggesting a recent common ancestry with minimal divergence. High bootstrap values across the phylogenetic tree underscore the low variability among these isolates. This high conservation level highlights the stable nature of the ToBRFV genome across diverse geographical regions and hosts (**Figure 1a**).

**Figure 1.**
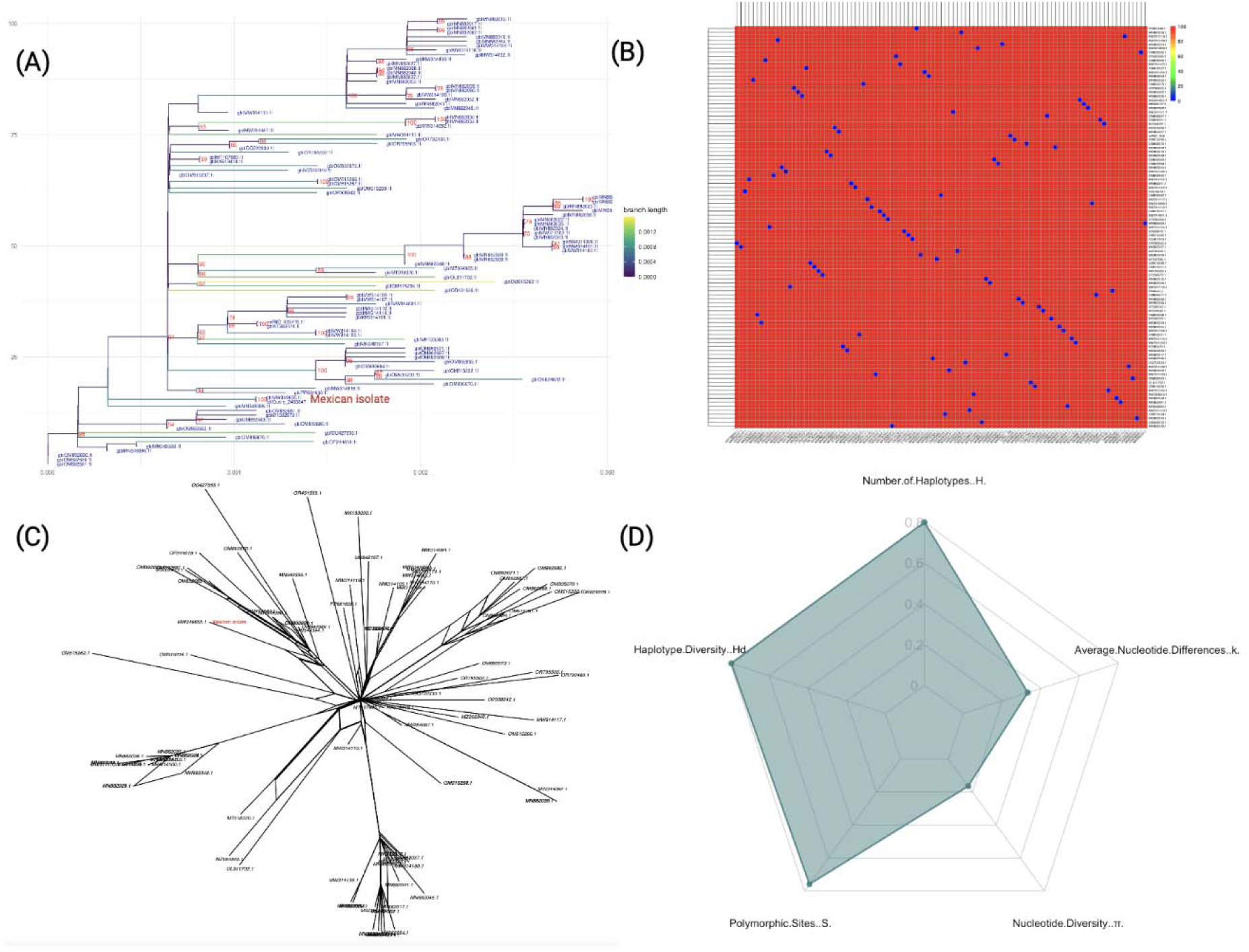
Comprehensive genetic diversity and phylogenetic analysis of ToBRFV isolates, highlighting the Mexican isolate. (A): Maximum-likelihood phylogenetic tree with 1000 bootstrap replicates, generated in `IQ-TREE2`. The Mexican isolate is highlighted in red, emphasizing its relationship to other isolates. Branch lengths are color-coded by distance to reflect sequence divergence. (B): Pairwise sequence identity heatmap for ToBRFV isolates, where colors represent nucleotide identity percentages, with warmer colors indicating higher conservation. (C): Neighbor- Net phylogenetic network depicting relationships among ToBRFV isolates, with the Mexican isolate labeled in red. This network visualizes potential lineage splits and diversification events. (D): Radar plot displaying genetic diversity parameters across isolates, including haplotype diversity (Hd), number of haplotypes (H), polymorphic sites (S), nucleotide diversity (π), and average nucleotide differences (k), illustrating the high genetic diversity and core conservation of the virus.

The Mexican isolate showed close genetic relationships with five other isolates, including three collected within Mexico, pointing to recent divergence events and supporting the possibility of local transmission. Notably, the Mexican TBRFV-MX-CP isolate from *Capsicum annuum* (2020) and isolates S33, S32, and S31 from *Solanum lycopersicum* (2018) suggest that greenhouse environments may serve as hotspots for ToBRFV spread. The phylogenetic proximity between the Mexican isolate and an Israeli isolate from *Solanum lycopersicum* seeds (2021) further suggests international transmission, likely facilitated by seed trade, underscoring the importance of stringent monitoring of seed imports to control viral spread. Collectively, these findings highlight a pattern of localized transmission within Mexico and the potential for global spread through international seed exchange, emphasizing the role of agricultural practices in the virus’s dissemination.

A detailed genetic diversity analysis further contextualizes the Mexican isolate evolutionary position (**Figure 1b**). Pairwise sequence identity calculations show strong nucleotide conservation across the 100 ToBRFV isolates, with the Mexican isolate displaying high sequence similarity to other isolates from Mexico and Israel. This high conservation level, with an average sequence identity of ∼99.79%, suggests minimal genetic divergence across isolates, reinforcing the stability of the ToBRFV genome across diverse locations and hosts.

To examine relationships more flexibly, we generated a Neighbor-Net phylogenetic network (**Figure 1c**). The limited branching in this network suggests a pattern of localized transmission within Mexico and supports the hypothesis of international spread through seed trade. This network view aligns with the high sequence identity findings, highlighting a viral population that maintains stability while adapting to new environments with only minor genetic variations.

Finally, genetic diversity metrics (**Figure 1d**) provide further insights, showing a pattern of high haplotype diversity (*Hd = 0.994*) with a significant number of unique haplotypes (*H = 80*) among the isolates. This high haplotype diversity indicates substantial genetic variation within the population. Despite this, both nucleotide diversity (π *= 0.0021*) and average nucleotide differences (*k = 0.2079*) remain low, suggesting that sequence differences are generally subtle and the ToBRFV genome remains highly conserved. The observed combination of high haplotype diversity and low nucleotide diversity reflects a population shaped by recent evolutionary and demographic events. This pattern suggests a history of rapid demographic expansion following a population bottleneck, where mutations accumulated in closely related haplotypes. Alternatively, purifying selection may have removed deleterious mutations, constraining nucleotide variability while allowing the proliferation of distinct haplotypes.

In summary, our findings suggest that ToBRFV, including the Mexican isolate, has evolved a balanced strategy of adaptability and stability, with sufficient genetic diversity to respond to environmental pressures while maintaining a highly conserved genome. This evolutionary pattern, likely driven by stabilizing selection, allows the virus to persist and spread effectively across regions.

### 3.2 Geographic and Host-Specific Adaptation in the Mexican ToBRFV Isolate

The geographical clustering of ToBRFV isolates highlights regional patterns that suggest localized adaptation and potential evolutionary divergence. Clades formed by isolates from distinct regions, such as Israel and Europe, point to regional evolutionary pressures shaping their genetic profiles. Within this context, the Mexican isolate exhibits a unique genetic position, balancing distinctiveness and similarities with geographically distant isolates, likely due to cross-regional transmission events. The distribution of genetically aligned isolates (**Figure 2A**) reveals strong affinities with the Mexican strain in regions such as Israel, China, and Mexico, while lower alignment frequencies in areas like the United States, Jordan, and the United Kingdom suggest varied evolutionary pressures and degrees of relatedness.

**Figure 2.**
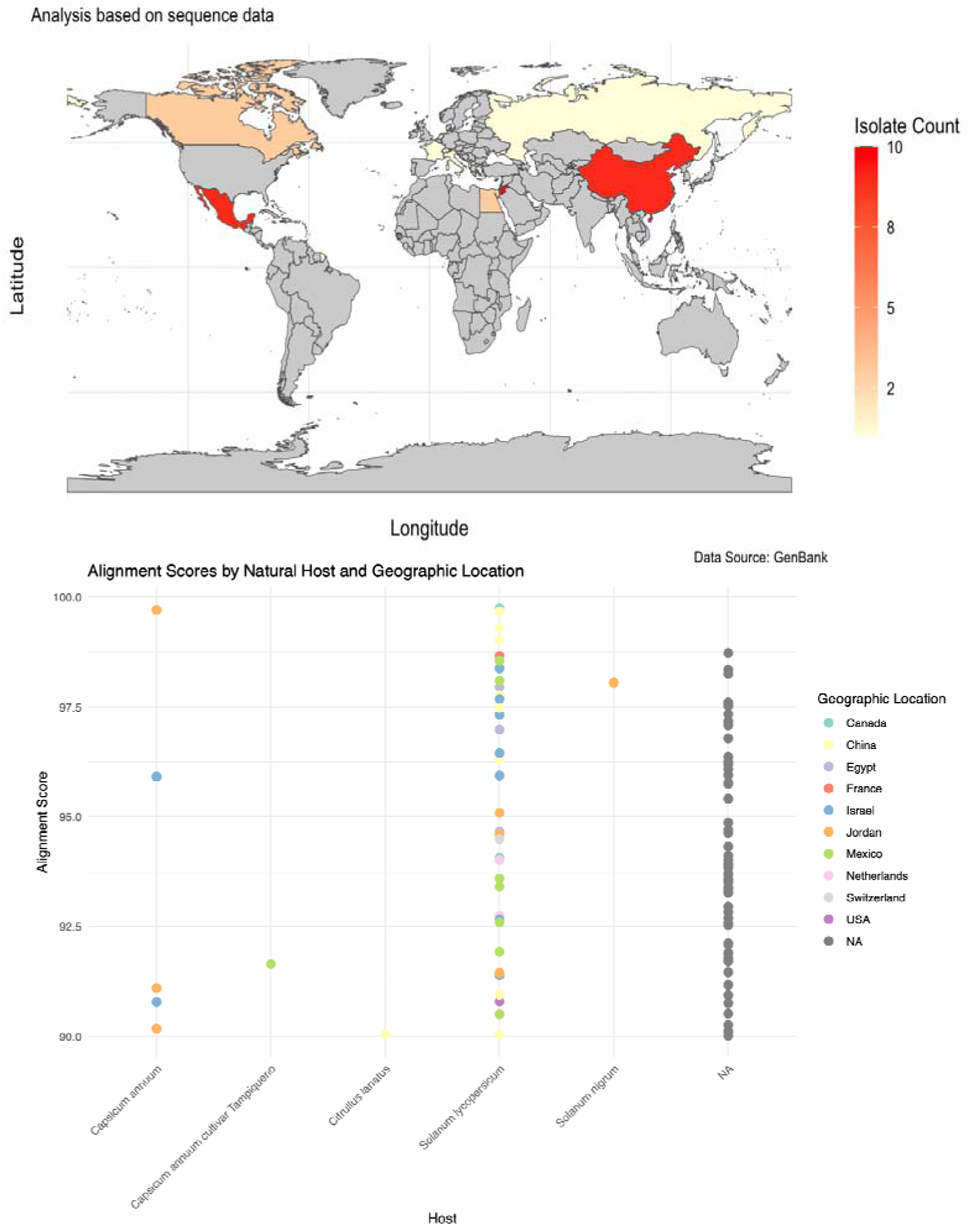
Alignment Scores of ToBRFV Isolate Geographic Location and natural host. **(A)**Geographical distribution map showing the number of ToBRFV isolates from various regions that align with the assembled genome of the Mexican strain. High isolate counts are observed in China and Mexico, with additional aligned isolates in the United States, Jordan, and the United Kingdom. The distribution highlights the broad geographical spread of genetically similar isolates and suggests potential routes of ToBRFV transmission, possibly facilitated by global agricultural trade. Data source: NCBI GenBank. **(B)** This figure displays alignment scores of ToBRFV isolates across various hosts and geographic regions. Each point represents an isolate, color-coded by geographic location. The alignment scores show a high degree of genetic similarity in isolates derived from *Solanum lycopersicum* (tomato), the host of the Mexican reference isolate. Lower scores in other hosts, such as *Citrullus lanatus* (watermelon) and *Capsicum annum* (pepper), indicate greater genetic divergence, suggesting host-specific adaptation and variability. Geographic analysis further highlights previously identified regional divergence among ToBRFV isolates. Higher alignment scores were observed in isolates from Mexico and China, indicating strong genetic similarity with the Mexican tomato-derived isolate.

Host-specific analyses provided additional insights into ToBRFV’s genetic variability and adaptive mechanisms (**Figure 2B**). Isolates from *Solanum lycopersicum* (tomato), the host of the Mexican reference genome, displayed high alignment scores, reflecting stable adaptation and a conserved genetic profile within this primary host. This suggests a recent shared ancestry between the Mexican isolate and other tomato-derived isolates.

In contrast, isolates from *Capsicum annuum* (pepper), and *Capsicum annuum* “*tampiqueño*” variety, showed moderate alignment scores, indicative of genetic divergence driven by host-specific selective pressures. Ortiz et al. (2022) demonstrated that the Mexican isolate caused the highest symptom severity in the *“tampiqueño”* chili variety among the eight tested, including necrotic lesions, chlorosis, mosaic, mottling, and leaf deformation. These results support the hypothesis that host-specific selective pressures influence virus-host interactions, as “*tampiqueño*” exhibited significant morphological and fruit quality impacts compared to less affected varieties, such as habanero.

In our analysis, greater variability was observed in *Citrullus lanatus* (watermelon) isolates, suggesting evolutionary adaptations that enhance ToBRFV fitness in non-tomato hosts. These patterns of alignment scores by host and geographic location (**Figure 2B**) highlight the virus’s ability to undergo distinct evolutionary trajectories in response to different host environments and defense mechanisms, underscoring the complexity of its adaptation and the need for host-specific management strategies.

### 3.3 Genomic Divergence in ToBRFV Replicase Regions: Insights into Host-Driven Adaptation

To gain deeper insights into the genomic structure and evolutionary pressures acting on the ToBRFV Mexican isolate, we conducted a genome comparative analysis focusing on specific nucleotide positions that exhibited significant variation in contrast to the 100 ToBRFV genomes (**Figure 3**). This analysis revealed five key nucleotide positions with normalized differences, potentially indicating sites of evolutionary divergence within the viral genome. Notably, the most variable position was observed at nucleotide 528, where a normalized difference of 1.000 suggests complete divergence at this site among the isolates, highlighting it as a hotspot for potential adaptation. Similarly, positions 1267 and 1881 displayed high levels of variability, with normalized differences of 0.897 and 0.867, respectively. The region surrounding nucleotide positions 528 and 1267 corresponds to the genomic span of 280–1365, which encodes a viral methyltransferase domain (*pfam01660*). This domain is involved in RNA cap methylation, enhancing RNA stability and ensuring compatibility with the host’s translation machinery. Variability in these positions may impact the efficiency of these processes, potentially influencing viral fitness and adaptation.Variations within this protein-coding region could enhance the virus’s fitness by enabling more efficient replication or evasion of host defenses, particularly if ToBRFV is adapting to new hosts or undergoing selection in greenhouse settings with intensive agricultural practices. Such adaptations could contribute to the virus’s persistence and ability to infect a diverse range of host species.

**Figure 3.**
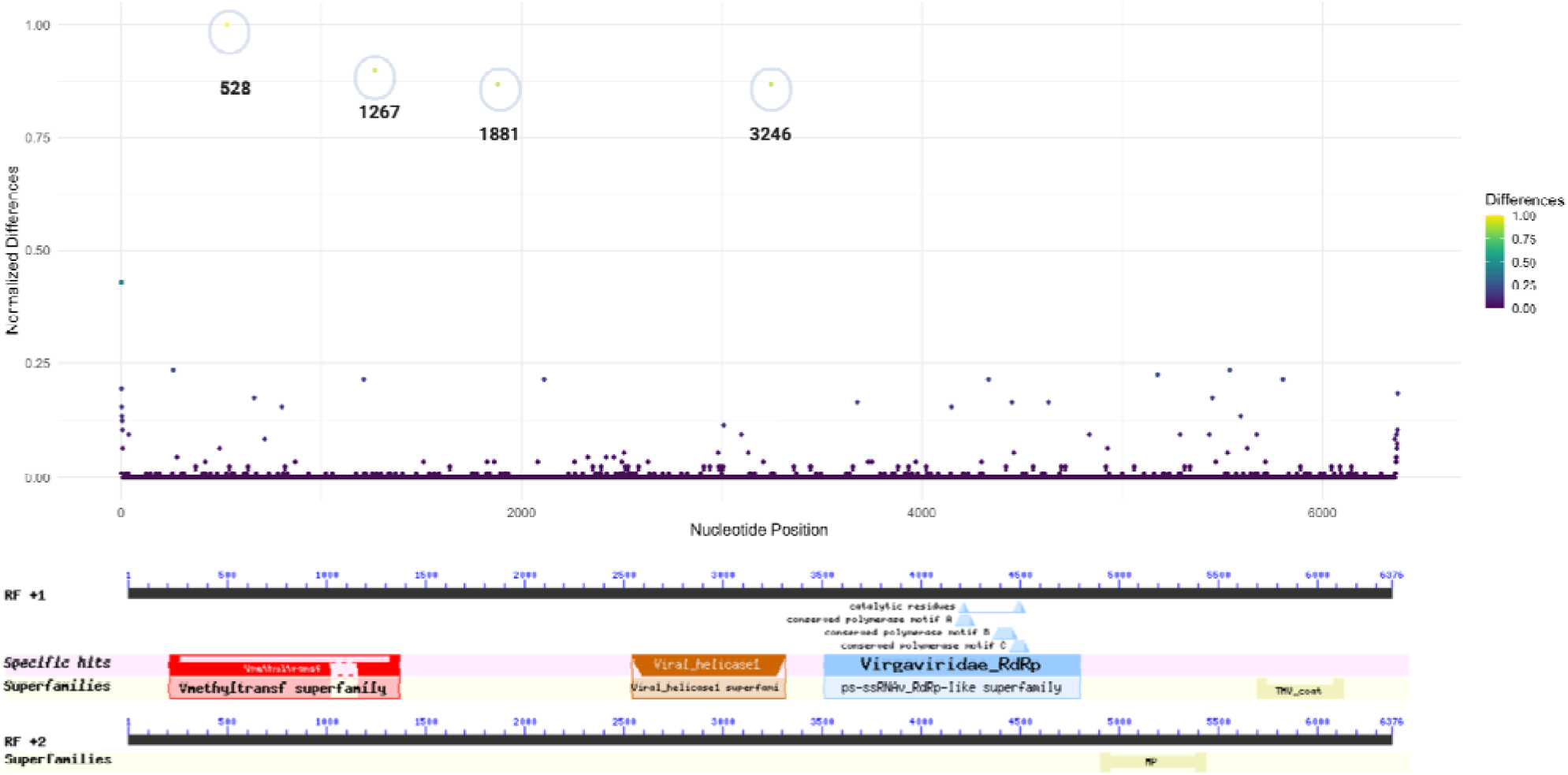
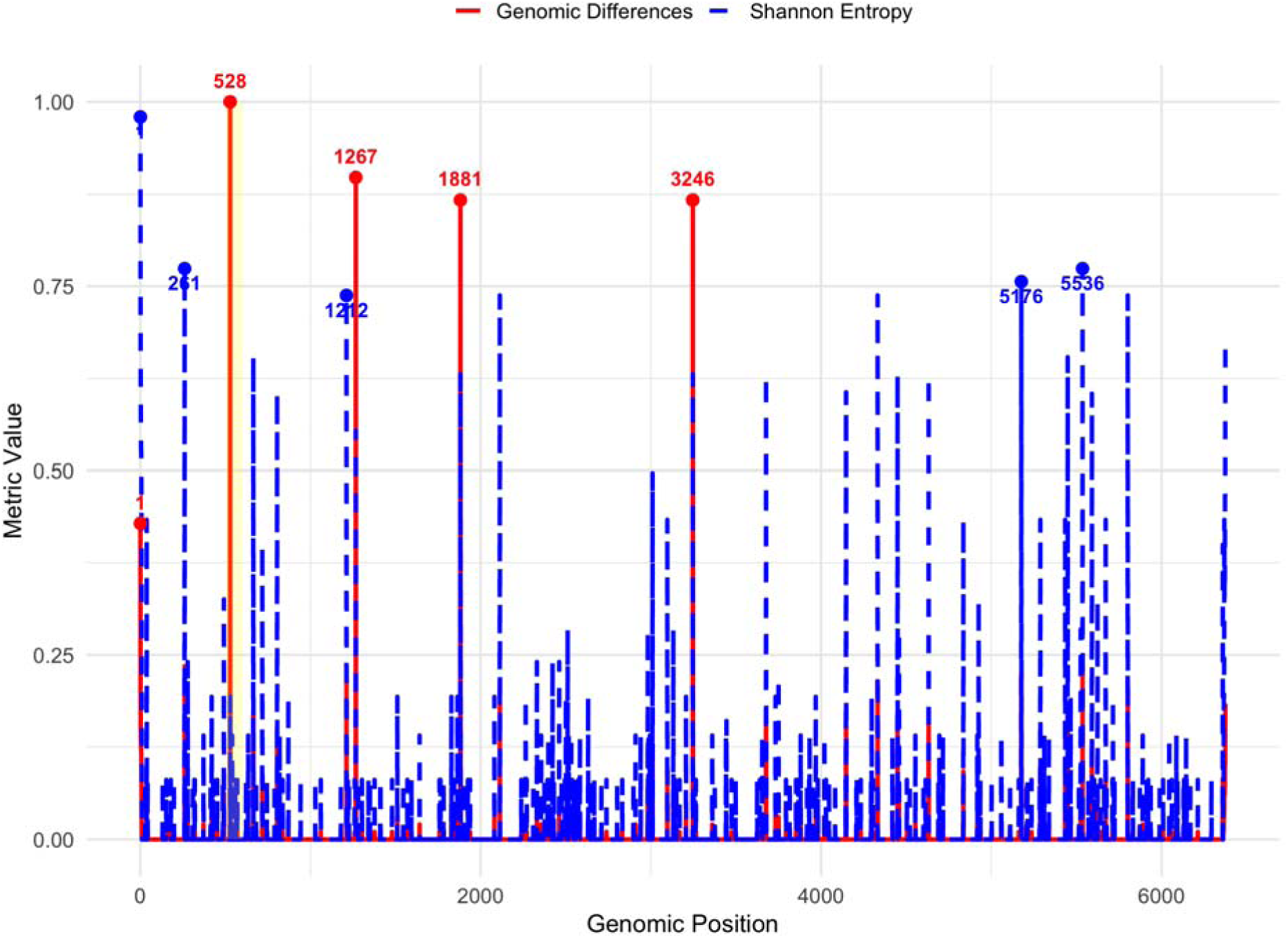
Genomic Differences of Mexican ToBRFV Isolate Compared to 100 Isolates. (A)This plot displays normalized nucleotide differences across the ToBRFV genome in comparison to 100 other isolates. Each point represents a nucleotide position, with the y-axis indicating normalized difference values and the x-axis showing nucleotide positions. Colors reflect the degree of variation, from low (purple) to high (yellow). Notable peaks at positions 528, 1267, and 1881 reveal regions of high genetic divergence, which correspond to genomic features associated with viral replication. (B) illustrates the specific functional domains identified within the ToBRFV genome. Superfamilies and conserved motifs are marked, demonstrating key structural and functional elements within the genome; (C)Comparison of genomic differences and Shannon entropy across the *ToBRFV* genome. The red solid line represents genomic differences, while the blue dashed line shows Shannon entropy. Key positions with the highest values for each metric are highlighted and annotated. A highlighted yellow region indicates an area of specific interest.

The region surrounding nucleotide position 1881 is not directly associated with any of the annotated functional domains in the current analysis. However, its high variability suggests it could represe t a previously uncharacterized functional element or a site under evolutionary pressure.

Nucleotide position 3246, with a normalized difference of 0.867, is located within the viral helicase domain (*pfam01443*, spanning 2543–3315, E-value: 4.31E-19), part of ORF1. This domain is essential for RNA unwinding during replication and transcription, highlighting its critical role in the viral life cycle. The observed variation at this position likely reflects selective pressures acting on the helicase function, potentially enhancing the virus’s adaptability and infectivity across diverse hosts and environments.

These findings indicate that the replicase coding regions, particularly ORF1, represent hotspots for genomic variation in the Mexican ToBRFV isolate. Variability within key domains, such as the methyltransferase and helicase domains, may play a pivotal role in the virus’s adaptation strategies, enabling it to optimize replication and transmission in diverse ecological contexts.

Shannon entropy analysis further revealed distinct variability patterns. A moderate positive Pearson correlation (0.66) and an almost perfect Spearman correlation (0.9999) between genomic differences and entropy suggest that higher sequence divergence aligns with greater sequence variability. Notably, position 1, with the highest Shannon entropy (0.979) and a significant genomic difference (0.429), emerges as a critical hotspot of genomic diversity with potential functional relevance.

While position 528 exhibits the highest genomic difference (1.000) but low entropy (0.193), indicating unique divergence specific to the Mexican isolate, positions with high entropy but low differences (e.g., 261, 5536) may represent conserved regions with broader alignment variability. Adaptive hotspots such as 1267, 1881, and 3246, characterized by both high differences (∼0.87) and moderate entropy (∼0.63), likely reflect regions under balanced evolutionary forces. These insights underscore the functional and evolutionary importance of replicase coding regions, particularly ORF1, in shaping the virus’s ecological adaptation and host interactions.

#### 3.3.1 Host-Specific Single Nucleotide Variability(SNVs) in Key Replicase Regions

The detailed nucleotide variability analysis reveals significant insights into the host-specific evolution and adaptability of ToBRFV. Isolates from *Solanum lycopersicum* exhibited highly conserved nucleotide profiles, particularly at positions V180, V414, V744, and V1377 within the replicase regions, **Figure 4**. This conservation underscores the virus’s genetic stability in its primary host and suggests a co-evolutionary relationship that has optimized ToBRFV’s replication and transmission capabilities in tomato. Such stability indicates that the virus has reached a near-equilibrium state in this host, reflecting selective pressures that maintain essential genetic features critical for fitness.

**Figure 4.**
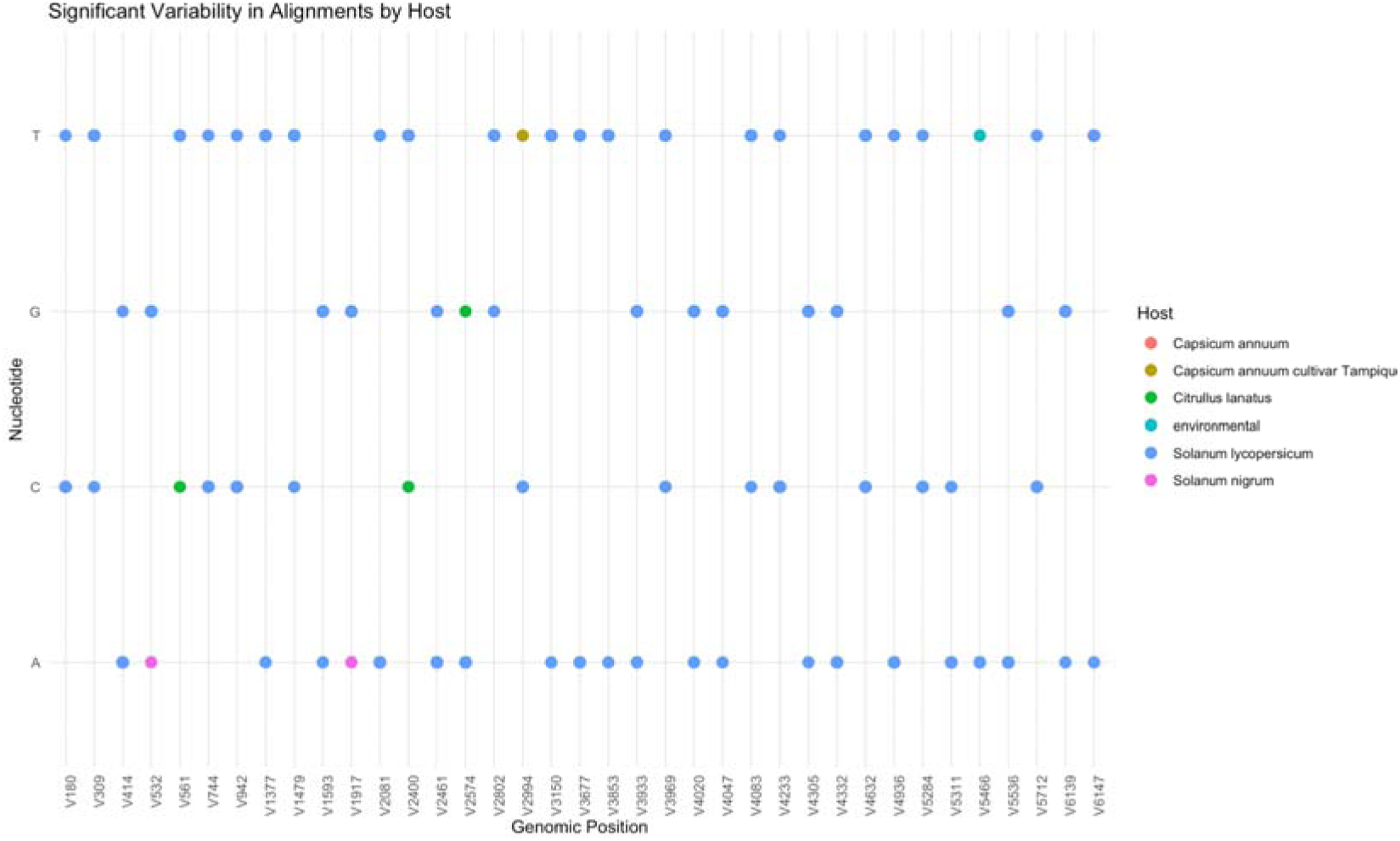
Significant Nucleotide Variability in ToBRFV Isolates by Host. This figure shows specific nucleotide positions in the ToBRFV genome where variability was observed across different hosts. Each dot represents a nucleotide position, color-coded by host species.

In contrast, isolates from alternative hosts, including *Capsicum annuum, Capsicum annuum* var. tampiqueño, and *Citrullus lanatus*, displayed unique single nucleotide variants (SNVs) that highlight host-specific selective pressures driving viral adaptation. For instance, substitutions at positions V532, V1377, and V2574 in *Capsicum annuum* suggest the virus is responding to distinct cellular environments, with *Capsicum annuum* var. tampiqueño showing slightly greater variability, likely due to localized ecological and molecular pressures. These findings indicate that ToBRFV has not yet achieved the same level of genetic optimization in pepper as observed in tomato, reflecting ongoing evolutionary divergence.

In *Citrullus lanatus* (watermelon), the identification of three notable SNVs at positions V561, V2400, and V2764 demonstrates a higher degree of genetic divergence. The substitution at position V561, located within the viral methyltransferase domain of the 126 kDa replicase protein, is of particular interest due to its potential impact on RNA cap methylation, a process critical for RNA stability and translation. The remaining substitutions fall outside annotated functional host-specific barriers in *Citrullus lanatus*. These substitutions may reflect evolutionary attempts by ToBRFV to establish infection in a less compatible host, highlighting the challenges the virus faces in adapting to non-tomato hosts. The greater variability observed in watermelon isolates underscores the incomplete adaptation process, which could involve modifications to critical genomic regions such as the replicase or other viral proteins.

Interestingly, *Solanum nigrum* isolates showed only two significant SNVs, both featuring adenine (A) at positions V52 and V917 within the coding region of the 126 kDa replicase protein. These positions overlap with the viral methyltransferase domain, suggesting a potential impact on replication efficiency. The limited variability in *Solanum nigrum* compared to other alternative hosts might reflect its role as a less favorable host, where selective pressures are weaker or less sustained due to limited viral replication or transmission success.

In *Capsicum annuum*, the thymine substitution observed at position V2994 within the helicase protein domain is particularly notable. These findings highlight the complex interplay between the virus and host-specific factors, driving genetic adaptations in non-primary hosts.

Overall, the presence of host-specific SNVs within the replicase regions underscores the remarkable genetic plasticity of ToBRFV. Variability within key functional domains, such as the methyltransferase, and helicase regions, likely confers selective advantages that enable the virus to fine-tune replication and host interactions. The strong genetic conservation among *Solanum lycopersicum* isolates reaffirms the tomato’s role as the primary host, where the virus has achieved a stable genomic configuration. In contrast, the higher nucleotide variability observed in non-tomato hosts, such as *Citrullus lanatus* and *Capsicum annuum*, reflects adaptive divergence driven by host- specific selective pressures.

All the mentioned genomic differences that occur in coding regions are likely non-synonymous, and could be altering the amino acids in the replicase proteins of ToBRFV. These SNVs may reflect adaptations to different hosts or, in some cases, incompatibility in cucurbit hosts. To assess their impact, we conducted an amino acid substitution analysis to evaluate how these mutations could affect the functions of the implicated domains (**Figure 5**).

**Figure 5.**
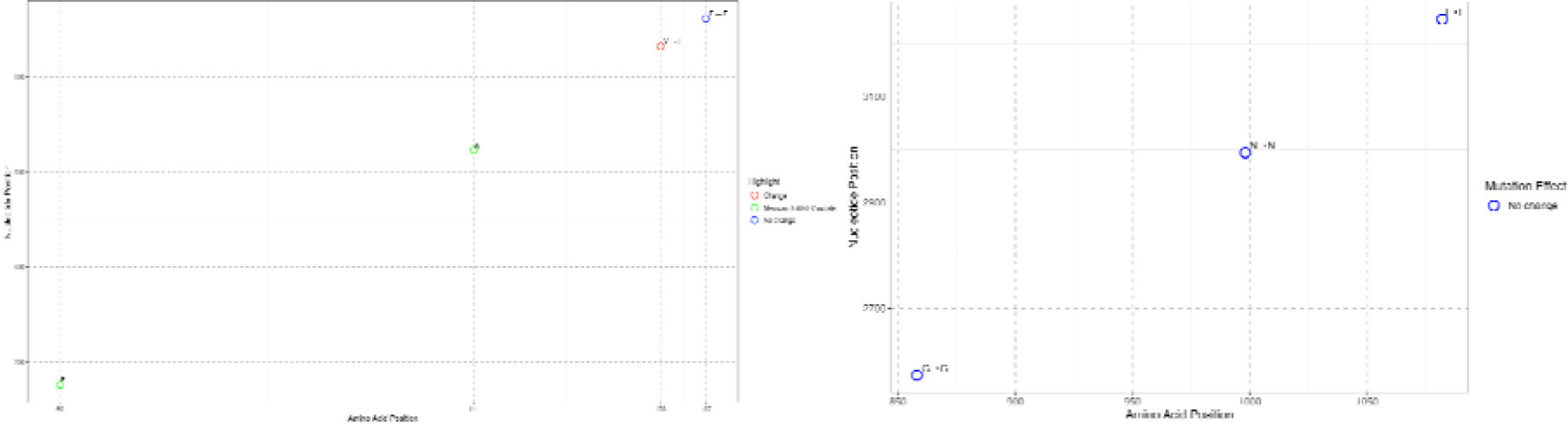
Amino acid substitution in ToBRFV replicase functional domains. **(A)** Methyltransferase domain: Substitutions at positions 59 and 141 are specific to the Mexican isolate, with position 178 (Valine-to-Isoleucine) linked to *Solanum nigrum*. Position 187 remains conserved in *Citrullus lanatus*.(B) Helicase domain: Conserved positions (e.g., 850, 1000, 3100) show no significant amino acid changes, maintaining structural and enzymatic stability.

The detailed amino acid substitution analysis, particularly within the methyltransferase domain of the replicase region, provides critical insights into the molecular adaptations of ToBRFV to its hosts and geographical distribution (**Figure 5A**). The results highlight the dual role of conserved and variable positions in maintaining functional stability while facilitating host-specific adaptability.

Amino acid positions 59 and 141, identified as the most divergent to the Mexican isolate, emphasize its geographic and genetic distinctiveness. These positions, which correspond to nucleotide changes within the methyltransferase domain, may represent adaptive modifications shaped by the ecological and agricultural context of Mexico. Position 59, in particular, resides within a region critical for RNA cap methylation, a key process required to stabilize viral RNA and ensure efficient translation. Such mutations may enhance replication efficiency in *Solanum lycopersicum*, reinforcing the evolutionary stability of ToBRFV in its primary host and enabling its continued prevalence in tomato populations in this region.

Position 178, characterized by a Valine-to-Isoleucine substitution, provides a compelling example of host-specific adaptation, particularly in *Solanum nigrum*. This mutation likely enhances viral infectivity by optimizing interactions with host-specific cellular machinery, supporting the hypothesis that ToBRFV undergoes selective pressures to overcome barriers in alternative hosts. Conversely, position 187 remains conserved as Phenylalanine in *Citrullus lanatus*, suggesting functional stability in this host. The lack of variability in *C. lanatus* indicates weak selective pressures, potentially reflecting limited evolutionary interaction or incomplete adaptation to cucurbit hosts.

The analysis further highlights conserved positions outside annotated functional domains, such as positions 850 and 3100, where no significant amino acid changes were detected (**Figure 5B**). These regions likely play structural or regulatory roles, maintaining the integrity of the replicase protein. Their conservation underscores their importance in ensuring the functional robustness of ToBRFV across diverse hosts. While these regions do not exhibit variability, they are essential for sustaining the core replicative machinery of the virus.

Overall, the observed patterns of variability and conservation within the replicase region reveal a strategic evolutionary balance employed by ToBRFV. Conserved residues within critical functional domains, such as the methyltransferase, ensure replication efficiency and stability in its primary host, *Solanum lycopersicum*. In contrast, variable residues identified in *Solanum nigrum* and *Citrullus lanatus* demonstrate the virus’s capacity for adaptive divergence in response to host-specific selective pressures. The close genetic similarity among Mexican *Solanum lycopersicum* isolates reflects the evolutionary stability of ToBRFV in tomato hosts, while the variability in non-tomato hosts points to adaptive divergence.

### 3.5 Expanded Host Susceptibility of ToBRFV in Native and Commercial Solanaceae

Following the *in silico* analysis of host specificity using available isolates from GenBank, experimental bioassays were performed to assess the susceptibility of various potential hosts to the Mexican ToBRFV isolate. The tested species included commercially cultivated plants—such as watermelon (*Citrullus lanatus*), cantaloupe (*Cucumis melo*), squash (*Cucurbita pepo*), cucumber (*Cucumis sativus*), pea (*Pisum sativum*), tomato (*Solanum lycopersicum*), and pepper (*Capsicum annuum*)—alongside less commonly cultivated or wild species, such as tomatillo (*Physalis ixocarpa*), tobacco (*Nicotiana rustica*), and eggplant (*Solanum melongena*).

The bioassays revealed strong host specificity for the Mexican ToBRFV isolate, with successful replication observed exclusively in solanaceous hosts, specifically tomato and pepper. Remarkably, the virus failed to replicate in any cucurbit hosts despite multiple inoculation attempts. This lack of replication suggests that the Mexican isolate lacks specific genetic adaptations required to efficiently interact with the cellular machinery of cucurbit hosts.

As summarized in **Table 2**, our findings confirmed tomato and pepper as susceptible hosts for the Mexican ToBRFV isolate. In contrast, no symptoms or detectable viral presence were observed in any tested Cucurbitaceae species, including watermelon, cantaloupe, squash, and cucumber. These results align with prior reports indicating that cucurbit hosts do not support ToBRFV replication under similar experimental conditions (Zhi-yong et al., 2021; Sabra et al., 2022).

**Table 2.** Analysis by nested RT-PCR and ELISA of ToBRFV inoculated plants.

| Inoculated specie | Nested RT-PCR | ELISA |
| --- | --- | --- |
| Watermelon (*Citrullus lanatus*) | 0*, 0°, 0" | 0, 0, 0 |
| Cantaloupe (*Cucumis melo*) | 0, 0, 0 | 0, 0, 0 |
| Squash (*Cucurbita pepo*) | 0, 0, 0 | 0, 0, 0 |
| Cucumber (*Cucumis sativus*) | 0, 0, 0 | 0, 0, 0 |
| Pea (*Pisum sativum*) | 0, 0, 0 | 0, 0, 0 |
| Tomato (*Solanum lycopersicum*) | 5, 5, 5 | 5, 5, 5 |
| Tomatillo (*Physalis ixocarpa*) | 3, 4, 3 | 3, 4, 3 |
| Tobacco (*Nicotiana rustica*) | 5, 5, 5 | 5, 5, 5 |
| Pepper (*Capsicum annum*) | 2, 4, 5 | 2, 4, 5 |
| Eggplant (*Solanum melongena*) | 0, 1, 5 | 0, 1, 5 |

Number of positive plants in the first (\*), second (°) and third (") experimental repetition. All the mock-inoculated plants were negative. Plants were analyzed after 25 dpi.

In contrast, experimental testing revealed new susceptibility insights for *Nicotiana rustica* (tobacco), *Physalis ixocarpa* (tomatillo), and *Solanum melongena* (eggplant). Interestingly, tomatillo and eggplant displayed partial infection, suggesting a degree of susceptibility not previously recorded. This variability in infection response highlights the nuanced host-pathogen dynamics within the Solanaceae family, with some individuals within species demonstrating possible resistance or lower susceptibility to ToBRFV. The symptomatic responses observed in infected plants, detailed in **Figure 6**, reflect these differences. Tomato and tobacco displayed classic ToBRFV symptoms, such as leaf narrowing and mosaic patterns, while tomatillo showed narrowing and chlorotic lesions, pepper exhibited stunting, and eggplant showed necrosis on both stem and leaves. In contrast, mock- inoculated plants (**Figure 6**, panels **6A-6E**) showed no symptoms, underscoring the specificity of ToBRFV-induced symptoms across susceptible hosts. These findings underscore the value of experimental validation to refine our understanding of the ToBRFV host range, especially in endemic and agriculturally relevant species. The observed variability in susceptibility across different species also highlights the complexity of ToBRFV’s host interactions and suggests the need for further research into the genetic and environmental factors influencing host susceptibility and resistance.

**Figure 6.**
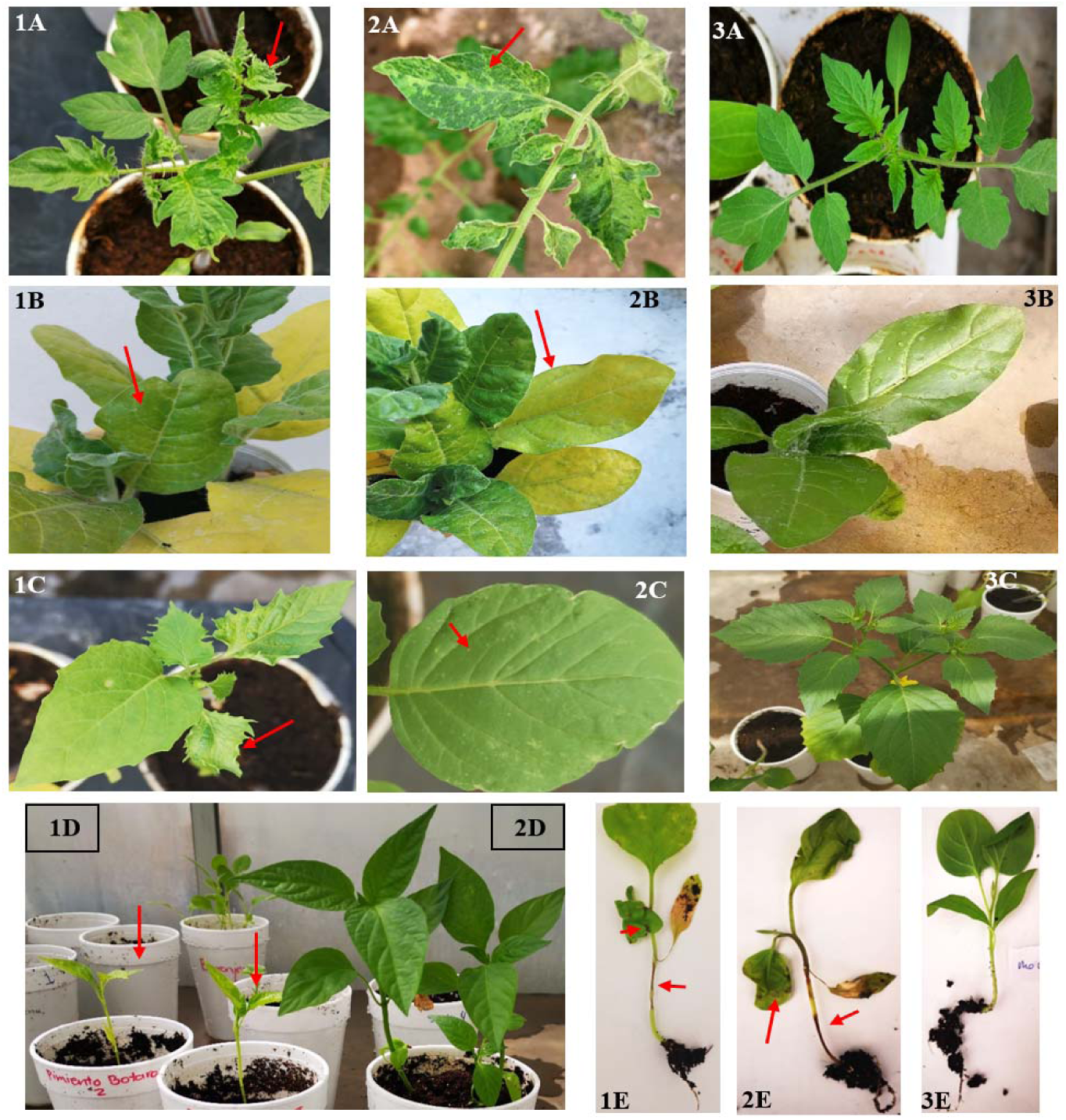
Symptoms in tomato (1-2A), tobacco (1-2B), tomatillo (1-2C), pepper (1D), and eggplant (1-2 E) infected with ToBRFV after mechanical transmission. Plants showed narrowing (1A, 1C), mosaic (2A), necrosis on stem and leaves (1E, 2E), chlorosis (1B, 2B), chlorotic lesions (2C), and stunting (1D). Mock-inoculated (3A,3B, 3C, 2D and 3E).

#### 3.6 Seed-Borne Transmission Dynamics

Following the host-specificity and infection susceptibility assessments, further investigations were conducted to evaluate the potential for seed-borne transmission in tobacco. This species was chosen due to its status as a plant with known susceptibility to ToBRFV and its ease of handling in controlled bioassays. These analyses enabled us to integrate genomic insights with biological evaluations of transmission and infectivity, constructing a comprehensive view of the behavior of the ToBRFV Mexican isolate across host interactions and transmission pathways.

To examine the effect of ToBRFV infection on seed viability, we analyzed germination rates in seeds from ToBRFV-infected and control *N. rustica* plants. The results demonstrated a significant reduction in germination rate for seeds from infected plants, with a mean germination rate of 57.7% compared to 82% for control seeds (**Table 3**).

**Table 3.** Comparison of Germination Rates: *N. rustica* Seeds from Healthy versus ToBRFV-Infected plants.

| Replicate | Number of germinated seeds |  |
| --- | --- | --- |
|  | ToBRFV-infected | Healthy-Control |
| 1 | 58 | 92 |
| 2 | 56 | 87 |
| 3 | 64 | 88 |
| 4 | 56 | 75 |
| 5 | 74 | 88 |
| 6 | 52 | 82 |
| 7 | 46 | 44 |
| 8 | 63 | 89 |
| 9 | 51 | 93 |

This decrease suggests that ToBRFV infection adversely affects the reproductive success of *N. rustica* by reducing seed viability. This finding aligns with earlier analyses suggesting complex host-pathogen interactions where viral infections impact plant reproduction beyond visible symptoms. The reduction in seed viability for infected plants has critical implications for agricultural biosecurity, particularly for preventing ToBRFV’s spread via seed trade.

#### 3.6.1 ToBRFV Infection Reduces Seed Viability and Enables Potential Seed-Borne Transmission in wild species, Posing Biosecurity Risks

To quantify the prevalence of ToBRFV within *N. rustica* seeds and seedlings, nested RT-PCR was conducted on varying subsample sizes, offering insights into virus detection variability. The infection rate in 150-seed subsamples was 0.61%, while 100-seed subsamples showed a lower infection rate of 0.22% (**Figure 7A**). Furthermore, ToBRFV presence was detected only in seedlings from the 150- seed subsamples, while no virus was found in seeds treated with a 3% sodium hypochlorite solution. These findings underscore the role of sample size and disinfection treatment in virus detection, highlighting sodium hypochlorite’s efficacy in removing surface-bound virions. However, the persistence of ToBRFV within internal seed tissues suggests that surface disinfection alone may be insufficient, emphasizing the need for rigorous testing protocols in seed transmission studies.

**Figure 7.**
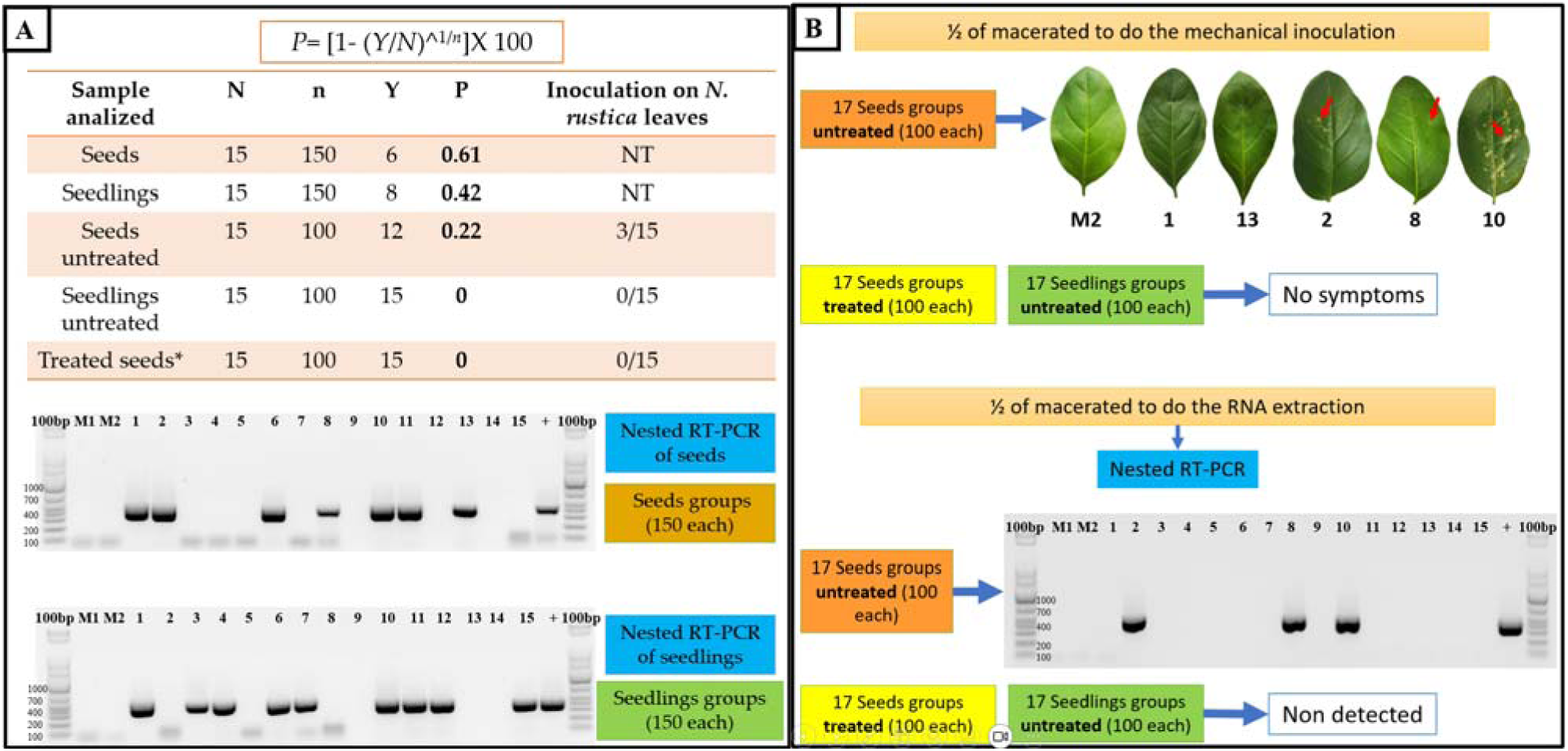
Seed-Borne Transmission and Infectivity Analysis of ToBRFV in *Nicotiana rustica*. **(A):** Seed transmission dynamics were evaluated through germination and nested RT-PCR analyses on *N. rustica* seeds and seedlings from ToBRFV-infected plants. Seed subsets with and without 3% sodium hypochlorite treatment were tested, and infection rates (P) were calculated using the formula: P=[1−(Y/N)1/n]P=[1−(Y/N)1/n], where NN is the total number of subgroups, nn is the number of seeds/seedlings per subgroup, and YY represents RT- PCR-negative subgroups. The electrophoretic gel image displays RT-PCR products (400 bp) for ToBRFV detection in infected and uninfected (control) samples, with M1 and M2 as healthy plant controls.**(B):** Viral infectivity in *N. rustica* was assessed by inoculating leaves with seed macerates from infected plants. Chlorotic lesions (arrow) indicate symptomatic infection, confirmed by RT-PCR (400 bp band). Treated seeds (sodium hypochlorite) showed no RT-PCR detection or symptoms, underscoring the role of surface disinfection in reducing infectivity.

#### 3.6.2 Assessing Seed-Borne Viral Infectivity

To further explore the infectivity potential of ToBRFV within *N. rustica* seeds, bioassays were conducted to observe symptom expression and confirm viral presence in inoculated plants. These bioassays revealed that only three groups of inoculated leaves displayed visible symptoms, which were confirmed as ToBRFV-positive through nested RT-PCR (**Figure 7B**). Conversely, no symptoms or virus detections were observed in seedlings derived from seeds treated with sodium hypochlorite, underscoring that viable ToBRFV particles may persist in untreated seeds, capable of initiating infection under favorable conditions. The absence of symptoms in certain cases further highlights the importance of molecular diagnostics for reliable virus detection, especially where symptoms may be latent or delayed.

The observed reduction in germination rate in ToBRFV-infected seeds raises several intriguing questions. While the phenomenon itself is clear – ToBRFV infection is associated with diminished seed viability – the underlying mechanisms are not yet fully understood. Viruses, particularly those of the tobamovirus family, are known to interact with host plants in intricate ways, potentially impacting various aspects of their physiology, including seed development. These findings bear significant implications, particularly in the context of agricultural biosecurity and seed trade. Reduced germination rates in infected seeds not only affect the plant’s reproductive success but can also lead to crop yield losses if such seeds are unknowingly sown. Furthermore, the precise mechanisms through which ToBRFV influences seed germination warrant further investigation, as this knowledge can inform strategies for mitigating the impact of the virus on crop production.

## 4 Discussion

The Mexican isolate of ToBRFV exemplifies a remarkable adaptability at both global and host-specific scales. This adaptability is evidenced by the virus’s genetic stability, environmental resilience, and its ability to infect a variety of plant hosts, spanning both commercial crops and endemic species. Consistent with other Tobamoviruses, ToBRFV’s capacity to infect a broad spectrum of Solanaceae species highlights its potential for persistence and spread across diverse ecosystems, especially in regions with rich agricultural and native plant diversity, such as Mexico. This adaptability not only underscores the virus’s potential impact on agricultural systems but also raises concerns about its spread and persistence in natural environments, where it could interact with a wider array of hosts and potentially drive new cycles of infection.

### 4.1 Global Transmission Potential Facilitated by Trade and Environmental Resilience

Our findings, in conjunction with recent studies, highlight the substantial global transmission potential of the Mexican ToBRFV isolate, pointing to a complex, interconnected network likely driven by international trade. In our genomic analysis, the Mexican isolate demonstrates significant genetic similarity to strains from China and Israel, underscoring its link to geographically distant regions. Recent evidence further supports this trend, with reports of a second ToBRFV introduction into Mexico involving a strain closely related to isolates from the Netherlands and the Middle East (Vargas-Mejía et al., 2023). This pattern suggests that ToBRFV may have entered Mexico through multiple, independent routes, facilitated by the global movement of agricultural commodities, particularly seeds and other plant materials.

The genetic proximity of the Mexican ToBRFV isolate to diverse international strains underscores its substantial potential for cross-regional movement, likely driven by trade-based pathways. This finding supports the hypothesis that ToBRFV’s global spread is facilitated through agricultural trade, enabling the virus to establish itself across continents. The presence of multiple introductions from distinct regions into a single country underscores ToBRFV’s capacity to traverse vast distances, effectively bypassing natural geographic barriers through the movement of infected plant materials and seeds. This mode of transmission not only highlights the adaptability of ToBRFV but also underscores the importance of implementing stringent biosecurity measures within trade networks to limit the virus’s continued spread and impact on agriculture worldwide.

The multiple independent introductions of ToBRFV from different geographic regions into Mexico’s agricultural systems raise the possibility of recombination events or regionally adaptive mutations. If genetically distinct strains co-occur within the same host populations, genetic exchanges could produce new ToBRFV variants with enhanced adaptability to local environmental and agricultural conditions. Such adaptive evolution could increase ToBRFV’s resilience and infectivity, complicating efforts to contain it in affected areas.This pattern of ToBRFV’s global movement, likely facilitated by trade networks, underscores an urgent need for coordinated international biosecurity initiatives. Monitoring seed imports and exports and enforcing rigorous quarantine protocols are essential strategies to limit the virus’s spread. These proactive measures can help mitigate ToBRFV’s impact on agriculture, especially in regions vulnerable to infection due to high trade volumes or diverse crop systems.

### 4.2 Genomic Stability and Host-Specific Adaptations of the Mexican *ToBRFV* isolate in Replicase domains

The genome of the Mexican ToBRFV isolate exemplifies a strategic balance between genetic stability and adaptive flexibility, allowing it to infect multiple hosts with minimal genetic drift. Comparative genomic analysis against 100 global ToBRFV isolates identified unique SNVs in the methyltransferase and helicase protein domains of the Mexican isolate, pointing to host-specific selective pressures. These protein domains are essential for viral replication, transcription, and immune suppression. Differences in these domains between the Mexican isolate and global genomes underscore their role in maintaining genetic integrity while facilitating viral persistence in both agricultural and natural ecosystems.

Our study corroborates recent findings that reveal ToBRFV’s low genetic variability and strong negative selection pressures, a pattern further supported by an analysis of an Iranian isolate (Esmaeilzadeh et al., 2023) which grouped ToBRFV into three clades with minimal genetic variation. This consistency across studies emphasizes the virus’s limited genomic diversity across geographic regions. Both studies demonstrate that ToBRFV is primarily subject to negative selection (dN/dS < 1), indicating an optimized viral genome for host infection with minimal adaptive pressure for mutation. This observation aligns with our findings of host-dependent single nucleotide variant (SNV) patterns within the replicase proteins, suggesting a stable genome that nevertheless supports host-specific adaptations.

The high genomic similarity observed in our Mexican ToBRFV isolate parallels findings by Van de Vossenberg et al. (2021), who reported 99.3% to 100% nucleotide identity across ToBRFV sequences from Dutch outbreaks, with up to 43 single nucleotide polymorphisms (SNPs) and limited amino acid changes. Notably, Van de Vossenberg et al. (2021) observed complete conservation in the 160 amino acid coat protein and minimal variability in the 1,621 amino acid RNA-dependent RNA polymerase (RdRp), with only up to six amino acid substitutions, underscoring the structural stability of these essential viral proteins. However, our study identifies subtle yet significant host-specific SNVs within the replicase protein, suggesting that, while ToBRFV maintains a stable genome, selective pressures may specifically influence this protein to support host-specific adaptations. This observation is further supported by Van de Vossenberg et al.’s findings of increased mutation rates in the movement protein (267 amino acids), which showed up to five amino acid changes, implying that certain functional domains may accommodate variability to facilitate viral spread across diverse hosts or environments. Specifically, the host-dependent variability in the methyltransferase and helicase domains identified in our study could confer adaptive flexibility, enhancing ToBRFV’s ability to establish infections across a range of hosts.

Recent studies also highlight the replicase domains as a focal point for adaptive flexibility, with this region showing the highest number of synonymous and non-synonymous mutations in ToBRFV (Abrahamian et al., 2022). The elevated mutation rate may confer a selective advantage, potentially enhancing ToBRFV’s interactions with diverse host factors, enabling infection across multiple host species. The presence of host-specific SNVs within this ORF in our study supports the protein’s role in host adaptation, particularly in *Solanum nigrum*, *Capsicum annuum*, and *Citrullus lanatus*. In *Capsicum annuum* and *Citrullus lanatus*, SNVs identified within the RdRp domains imply modifications that could affect replication efficiency within these hosts.Genomic variability analysis of the Mexican isolate revealed key SNVs in domains associated with replication, movement, and host interaction. For example, within *Capsicum annuum*, a unique thymine (T) substitution at position V2904, located in the 183 kDa replicase protein, suggests a host-specific adaptation that may enhance replication efficiency within pepper tissues. This mutation likely facilitates interaction with pepper-specific factors, contributing to the virus’s ability to establish infection in this solanaceous host.

Interestingly, the three distinct SNVs found in the Mexican ToBRFV isolate within *Citrullus lanatus* may limit viral replication in this non-solanaceous host, suggesting that specific nucleotide variations in the replicase regions may act as genetic barriers, reducing compatibility with *C. lanatus*’ cellular machinery. Structural or functional constraints in the viral replicase could hinder optimal interactions with host-specific factors, thus impeding successful replication. Future studies could target the methyltransferase and helicase domains of the Mexican ToBRFV isolate. Mutating SNVs to match sequences from compatible hosts could reveal their role in host restriction, particularly in *Citrullus lanatus*

In contrast, the absence of adaptive SNVs in regions critical for host compatibility in cucurbits, such as the helicase and methyltransferase domains, may explain the inability of the Mexican ToBRFV isolate to replicate in cucurbit hosts. This lack of host-specific adaptation could limit the isolate’s interaction with cucurbit cellular processes, preventing successful replication. Additionally, cucurbit hosts may exert different selective pressures that the Mexican isolate has not encountered or adapted to, suggesting a potential evolutionary divergence within ToBRFV populations that enhances host specificity.

Overall, our results demonstrate that while ToBRFV’s genomic stability supports high infectivity in its primary host, tomato, its broader adaptability allows it to infect diverse alternative hosts. This adaptability may enable ToBRFV to establish reservoirs in non-crop plants, promoting environmental persistence and transmission potential.

### 4.3 Expanded Host Range: Implications of Susceptibility in Endemic and Commercial Species

Building on the observed host adaptability, experimental bioassays have revealed susceptibility to Mexican ToBRFV isolate in *Nicotiana rustica*, *Physalis ixocarpa*, and *Solanum melongena*, further expanding the known host range of the virus. This extended host range reinforces ToBRFV’s potential to infect not only economically vital crops but also native plants, which could act as viral reservoirs within Mexico. Given the high transmissibility of ToBRFV, the infection of *Physalis ixocarpa* (tomatillo), a staple in Mexican agriculture, raises particular concern due to the potential for localized outbreaks that could threaten regional production.

In comparison with recent studies, our findings provide a targeted view of ToBRFV host adaptability, specifically in controlled greenhouse environments and seed-borne transmission contexts. While previous research has identified *Ipomoea purpurea*, *Mirabilis jalapa*, *Clematis drummondii*, and *Solanum tuberosum* as additional natural hosts in field settings (Eichmeier et al., 2023); our study expands the known host range by confirming susceptibility in *Physalis ixocarpa* and *Solanum melongena*. These findings suggest that ToBRFV exhibits diverse adaptability across solanaceous and non-solanaceous species, which could play a significant role in viral persistence in close proximity to tomato crops.

Moreover, findings have shown that ToBRFV has naturally infected eggplant in Mexico (EPPO-192, 2019). Panno et al. (2019) were not able to experimentally infect this vegetable species under controlled conditions with 14 hours of light at 20° to 28°C, while Fidan et al. (2020) did detected by RT-PCR to the ToBRFV in artificial inoculated eggplants but all were asymptomatic, using the same temperature but a photoperiod of 16 h of light. Variations in temperature and in the photoperiod can undoubtedly determine whether the virus successfully infects these plants and causes symptoms.

The susceptibility of *Solanum melongena* (eggplant) to systemic infection under specific experimental conditions aligns with findings that variations in temperature and photoperiod can influence virus infectivity (Szittya et al., 2003; Fajinmi and Fajinmi, 2010). Studies indicate that systemic infection rates and symptom expression in infected plants increase at higher temperatures, likely because these conditions promote viral replication and movement within the host (Chellappan et al., 2005; Zhang et al., 2012; Chung et al., 2015). This temperature-dependent infectivity suggests that environmental factors may play a significant role in shaping the epidemiology of ToBRFV, affecting infection dynamics across regions and climates. These findings underscore the need to consider ecological and climatic factors in managing the spread and impact of ToBRFV across different agro-ecological zones.

### 4.4. Seed-Borne Transmission and Biosecurity Risks in *Nicotiana rustica*

To gain deeper insights into the Mexican ToBRFV isolate adaptability and persistence, we conducted seed-borne transmission studies in *Nicotiana rustica*, which underscore the virus’s potential biosecurity risks. Infected seeds demonstrated a 30% reduction in germination rates compared to healthy controls, highlighting ToBRFV’s ability to compromise seed viability. A reduced percentage of germination has been documented in some plant’s infections with a viral complex (Marodin *et al*., 2019; Hemmati and McLean, 1977). Although the phenomenon that is involved is not well known, the viruses can clearly affect the physiological performance of the seed.

In tomato, the percentage of ToBRFV seed transmission has been recorded between 9% and 1.8% (Vargas-Mejía et al., 2023; Davino et al., 2020). The percentage of virus transmitted by the seed is different among viral species as well as among the variants of the same virus. This percentage can be modified by different factors, including: mixed infections, the type of host and the stage of development during which it is infected, the severity and environmental factors that affect both the host and the performance of the virus like its mobility and multiplication in the inflorescence (Mohan et al., 2020; Cobos et al., 2019). In addition, the viral accumulation in the host and the type of transmission (vectors, contact, seed) can modify the rate of transmission of the virus (Froissart et al., 2010; Pagán, 2023).

Despite disinfection treatments, the virus persisted within seeds, posing a significant risk given the global nature of seed trade. Though infection rates in seeds and seedlings were below 1%, even minimal transmission can enable the virus to establish infection sites far from its original source, reinforcing the urgent need for biosecurity protocols (Mohan et al., 2020). The combined challenges of host adaptability, environmental resilience, and seed-borne transmission make ToBRFV a formidable pathogen with global implications for agricultural trade and plant health.

This work detected the virus in subsamples of 150 and 100 seeds. According to Dombrovsky and Smith (2017), the threshold for detecting Tobamovirus in seeds is one infected out of 249, and 20 subsamples of 100 seeds are required to ensure a 95% likelihood of detecting the virus in minimal infestations of 0.15%. In the present work, we have observed some advantages and disadvantages that need to be analyzed to determine whether ToBRFV can be detected directly from the seed or the seedling. Seed samples can be immediately investigated, and a more significant number of seeds can be used for each sample processed during extraction, thereby increasing the likelihood of detecting the virus if a few seeds are infected. Nevertheless, non-viable virions, such as contaminants, can be found in the seed, which increases the risk of false positives. In addition, because of the seed tissue’s hardness and structural complexity, obtaining a high-quality RNA extraction is more complicated.

While using seedlings makes it possible to increase the replication of the virus and thereby increase the likelihood of detection, it has been found that seedlings remain asymptomatic (with low viral load) and do not express symptoms until handled in intensive production. Meanwhile, analyzing only seedlings without the seed coat eliminates the risk of false positives. The seedling tissue is easier to process, increasing the quality of the extracted RNA. Nonetheless, fewer can be processed since the plant material weighs more in seedlings. There needs to be more information about how many seedlings per subsample need to be processed to detect tobamoviruses.

In general, treating seeds with 1-3% sodium hypochlorite (NaOCl) has been shown to be effective against viruses (Herrera-Vásquez *et al*., 2009), and it has been corroborated that 2.5% NaOCl for 15 minutes was able to inactivate ToBRFV (Davino *et al*., 2020). The efficacy of sodium hypochlorite as a surface disinfectant was confirmed, as treated seeds showed no positive RT-PCR detection of ToBRFV. This finding has direct implications for seed certification protocols, as disinfection measures could effectively reduce the risk of ToBRFV transmission through contaminated seeds. However, the virus’s persistence within seed tissues despite disinfection highlights the need for molecular diagnostic tools capable of detecting internal infections. Incorporating these tools into routine biosecurity practices is essential to prevent ToBRFV’s spread via asymptomatic, infected seeds in international markets.

Our findings underscore the need for an integrated management approach to address the multifaceted nature of ToBRFV, particularly its host versatility, seed-borne transmission potential, and environmental resilience. The identification of host-specific adaptations within replicase proteins highlights the importance of developing resistant cultivars tailored to regional viral strains. In summary, the Mexican isolate of ToBRFV exemplifies the virus’s dual ability to maintain genomic stability while adapting to diverse hosts and environments, a characteristic that reinforces its position as a significant global pathogen. This study contributes valuable insights into the virus’s adaptability and suggests practical biosecurity and management strategies, ultimately supporting the global agricultural sector’s efforts to contain and mitigate ToBRFV’s impact.

## 5 Conclusions

Our study on the Mexican isolate of ToBRFV highlights the virus’s genetic variability, especially within the replicase regions. Additionally, the close genetic ties between the Mexican isolate and international strains point to global trade as a key factor in ToBRFV dissemination, underscoring the need for stringent biosecurity protocols in the seed trade. The potential role of native plants as viral reservoirs and the temperature-dependent infectivity observed in eggplant suggest that ToBRFV’s impact may also be climate-sensitive. In summary, ToBRFV’s genetic adaptability and transmission potential confirm its status as a global agricultural threat, calling for an integrated management approach involving breeding for resistance, enhanced biosecurity, and environmental monitoring.

## 3. Author Contributions

Conceptualization: ZMEJ and KAP. Data curation: KAP. Formal analysis: ZMEJ. Funding acquisition: HRW and OMDL. Investigation: CCCY and EJZM. Methodology: ZMEJ, KAP and CCCY; Formal Analysis, ZMEJ and OMDL; Project administration: ZMEJ and KAP. Resources: ZMEJ and OMDL. Software: KAP. Supervision: HRW and KAP. Validation: ZMEJ and KAP. Visualization: ZMEJ, KAP, HRW and OMDL. Writing – original draft: ZMEJ and KAP. Writing – review & editing: ZMEJ, KAP, HRW.

## 5. Acknowledgments

This is a short text to acknowledge the contributions of specific colleagues, institutions, or agencies that aided the efforts of the authors.

## 6. Supplementary Material

Supplementary Material should be uploaded separately on submission, if there are Supplementary Figures, please include the caption in the same file as the figure. Supplementary Material templates can be found in the Frontiers Word Templates file.

Please see the Supplementary Material section of the Author guidelines for details on the different file types accepted.

## 7. Data Availability Statement

The datasets generated and analyzed for this study can be found in the GitHub repository https://github.com/kap8416/TOBRFV-Genome-Analyses.

---

**Supplemental Figure 1.**
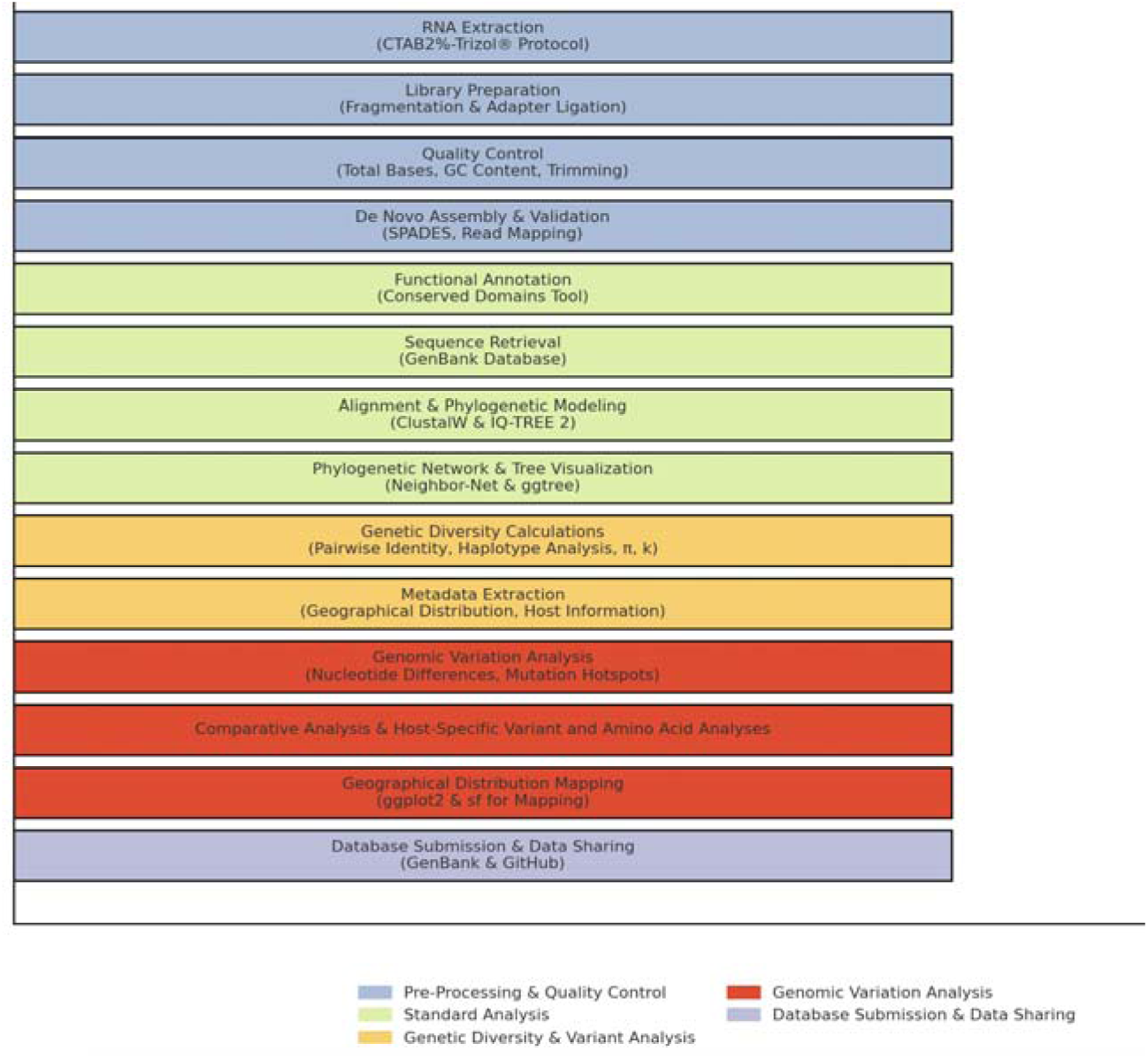
Workflow for the *in silico* analysis of the ToBRFV Mexican isol te. The workflow includes the following key steps, categorized by function: *Pre-Processing & Quality Control (blue)*: Includes RNA extraction, library preparation, quality control, and de novo assembly and validation. *Standard Analysis (green*): Functional annotation, sequence retrieval; full genome sequences of 100 ToBRFV isolates are retrieved from GenBank, alignment and phylogenetic modeling; sequences are aligned with `ClustalW`, and `IQ-TREE2` is used to perform phylogenetic modeling with 1000 bootstrap replicates, and phylogenetic network visualization; `Neighbor-Net` and `ggtree` in R are used to visualize the phylogenetic relationships among isolates. *Genetic Diversity & Variant Analysis (orange)*: Genetic diversity calculations (pairwise identity, haplotype analysis, π, k), metadata extraction geographical and host information is extracted for each isolate, and geographical distribution mapping. *Genomic Variation Analysis (red):* Nucleotide differences, mutation hotspot identification, and comparative host-specific variant and amino acid analyses. *Database Submission & Data Sharing (purple)*: Final step for submission and sharing of annotated data with GenBank and GitHub.

**Supplemental Figure 2.**
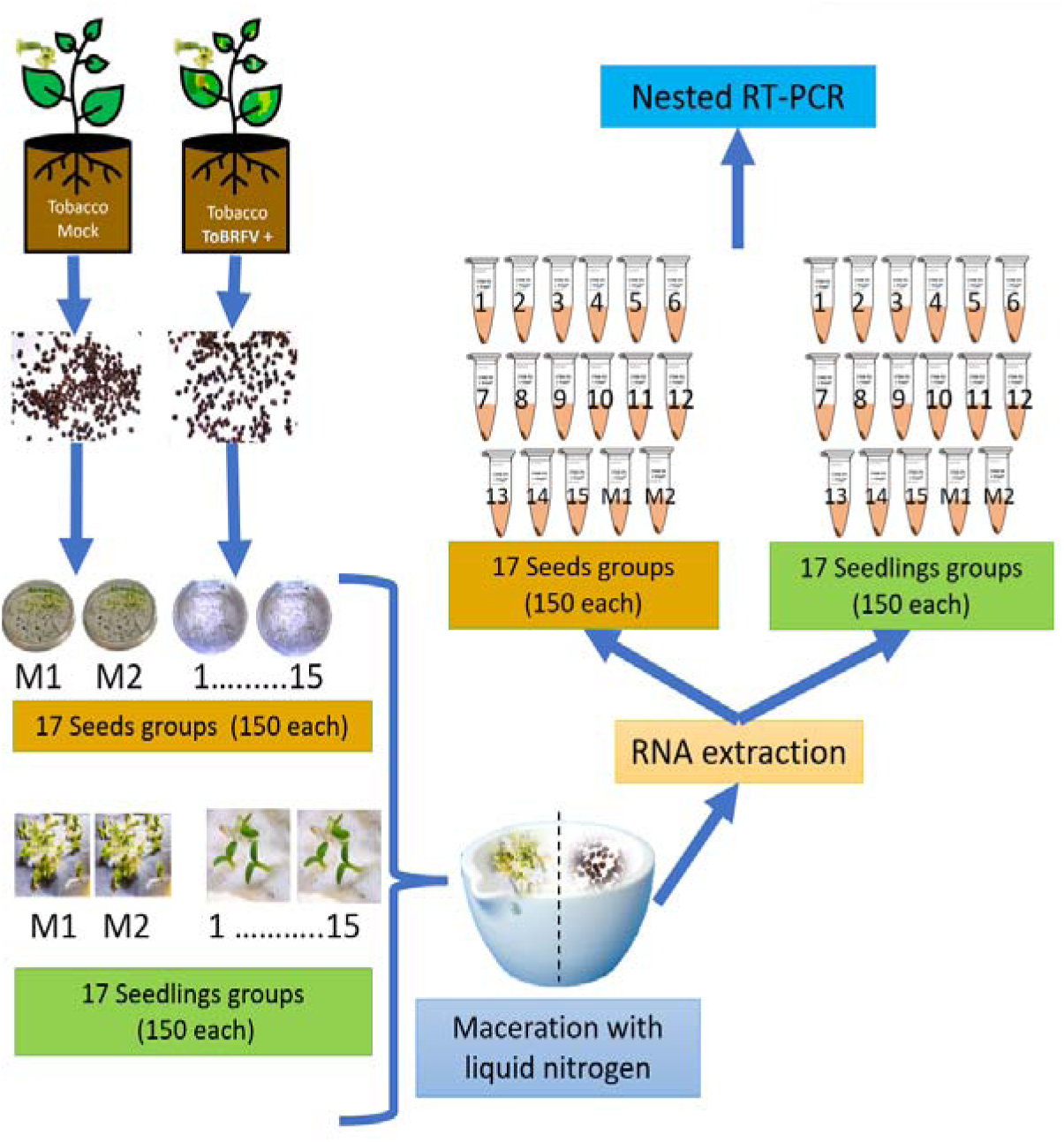
Schematic detailing the procedure for determining the percentage of infection in seeds and seedlings. Groups M1 and M2 symbolize mock-inoculated controls. numbers 1-15 represent individual groups of seeds and seedlings examined.

**Supplemental Figure 3.**
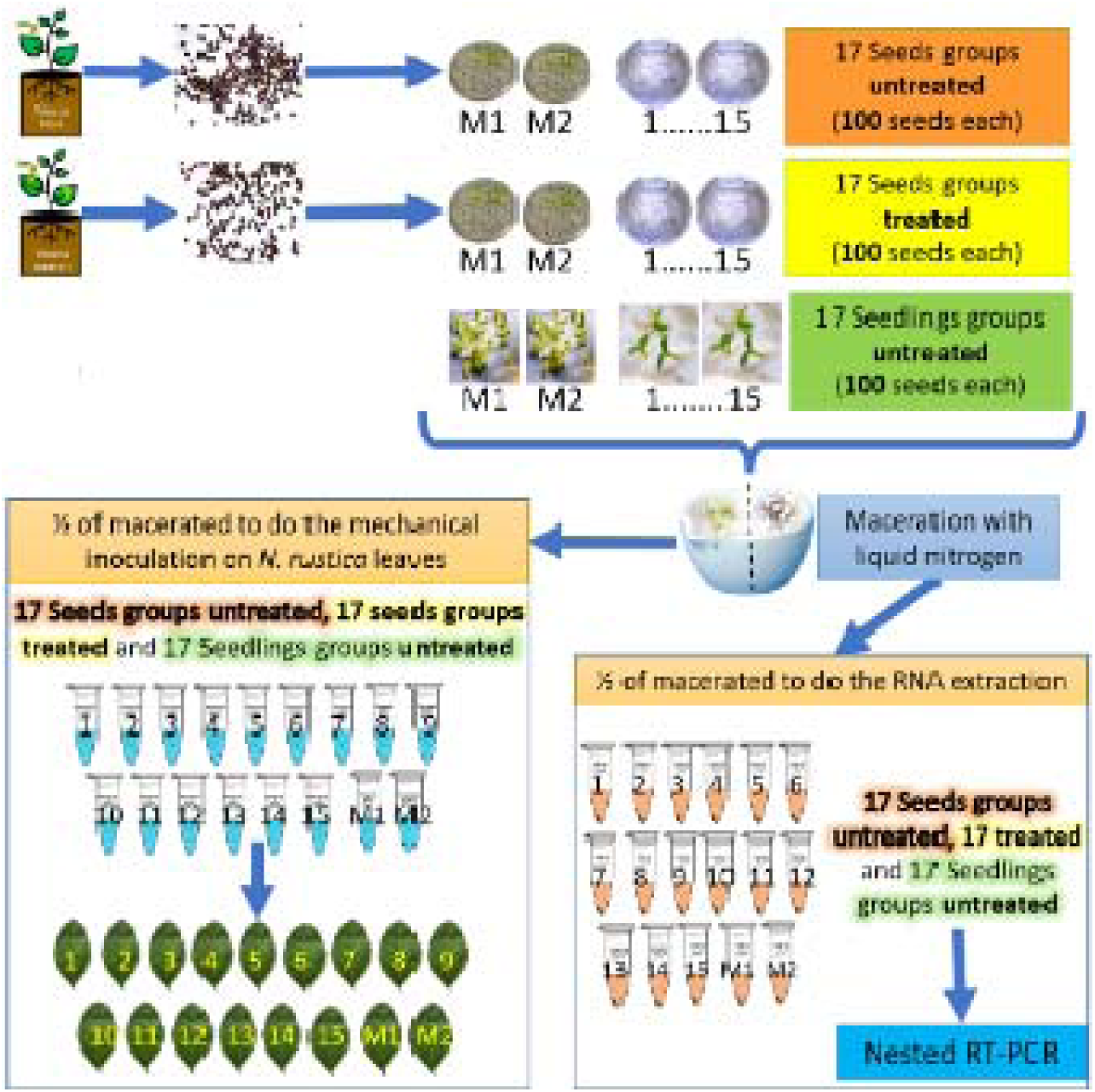
Flowchart illustrating the evaluation of viral particle infectivity in seeds. M1 and M2 denote the mock-inoculated controls. Groups 1-15 signify distinct batches of seeds and seedlings. Notably, all seed groups underwent a treatment with a 3% sodium hypochlorite solution for a span of 3 minutes.

**Supplemental Figure 4.**
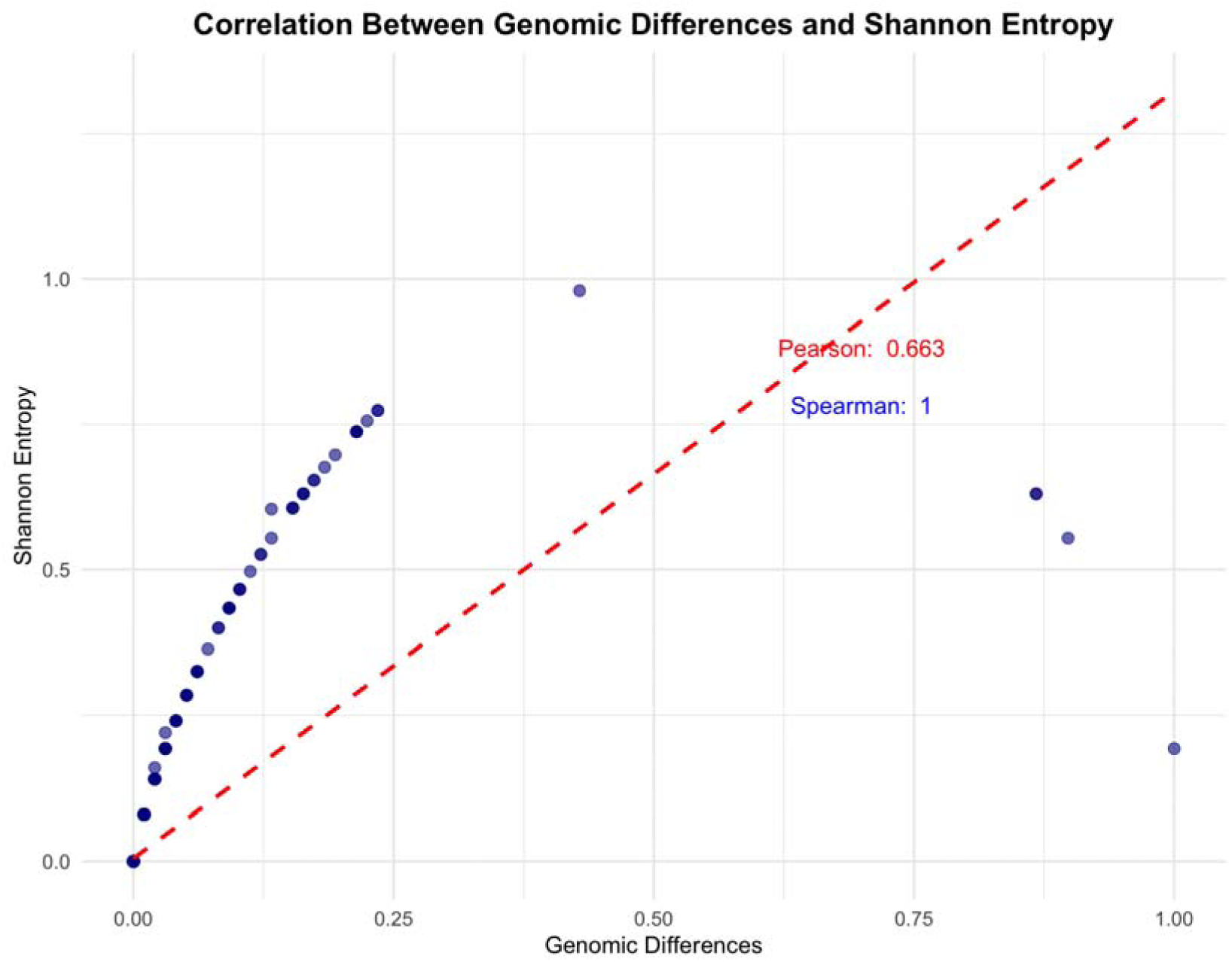
Correlation Between Genomic Differences and Shannon Entropy. Scatter plot showing the relationship between genomic differences (x-axis) and Shannon entropy (y- axis) for positions in the *ToBRFV* genome. The red dashed line represents the Pearson correlation coefficient (r=0.663), indicating a moderate linear relationship. The Spearman correlation coefficient (r= 0.999) highlights an almost perfect monotonic relationship, suggesting that positions with higher differences consistently show greater sequence variability.

## Supplemental material

1. Parameters of Mexican ToBRFV isolate sequencing

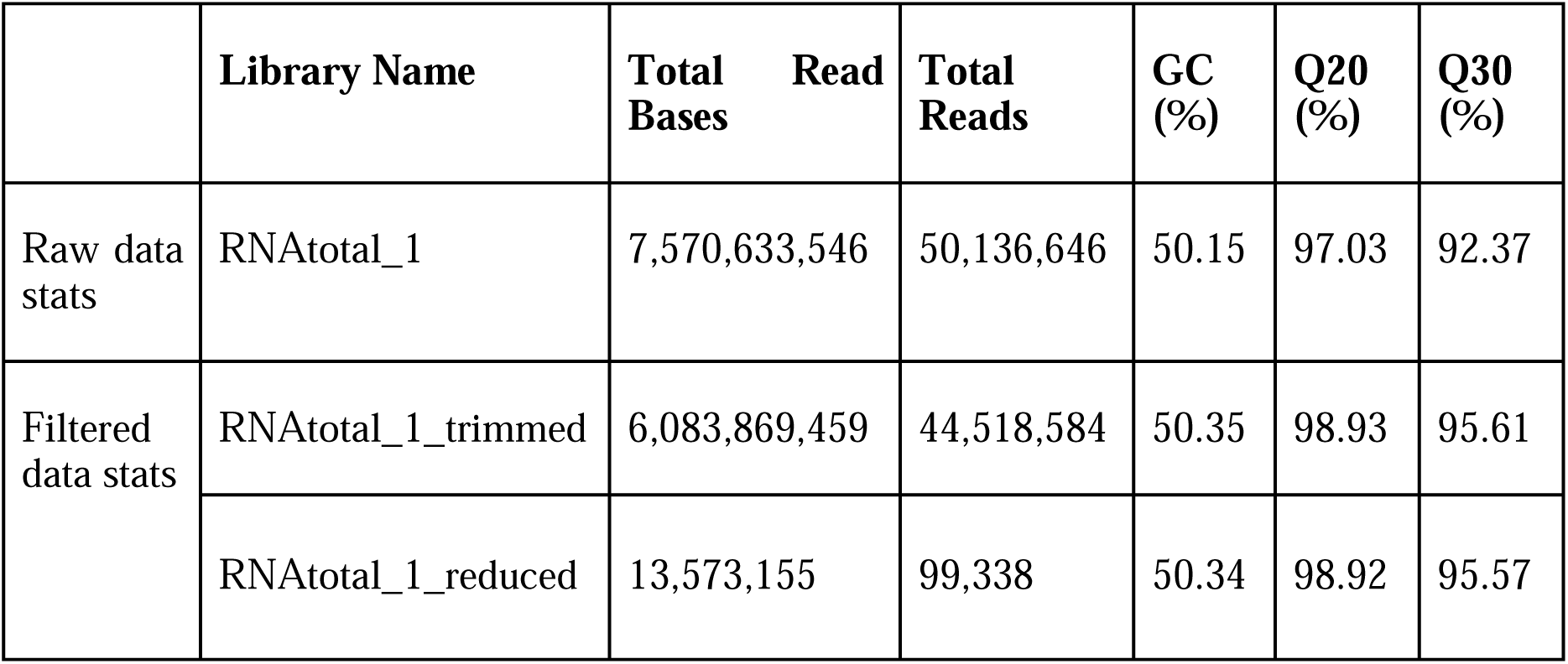
2. Base contents of contigs

3. Overall mapping stats

2. Base contents of contigs
| Num of A | Num of T | Num of G | Num of C | Num of N | GC contents |
| --- | --- | --- | --- | --- | --- |
| 1938 | 1789 | 1467 | 1181 |  | 41.5% |

4. Type of genome

3. Overall mapping stats
| Library Name | Total reads | Mapped reads | Coverage (%) | Depth | Ins. Size (Std.) |
| --- | --- | --- | --- | --- | --- |
| RNAtotal_1 | 99338 | 31201 (31.41%) | 100.00 | 607.16 | 171.62 (59.66) |
| Total | 99338 | 31201 (31.41%) | 100.00 | 607.16 | - |

4. Type of genome
| Contig name | Length | GC (%) | Circular |
| --- | --- | --- | --- |
| Contig1 | 6375 | 41.54 | NO |
| Total | 6375 | 41.54 |  |

**Supplemental Figure 5.**
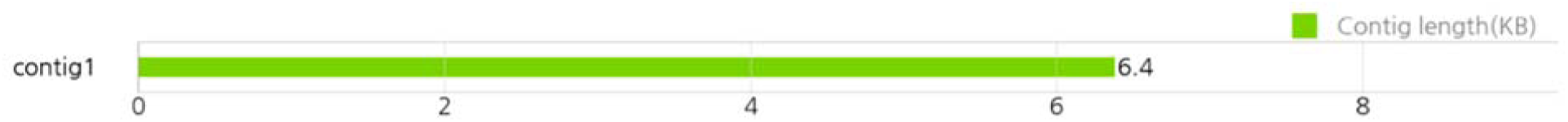
Figure of longest contigs.

**Supplemental Figure 6.**
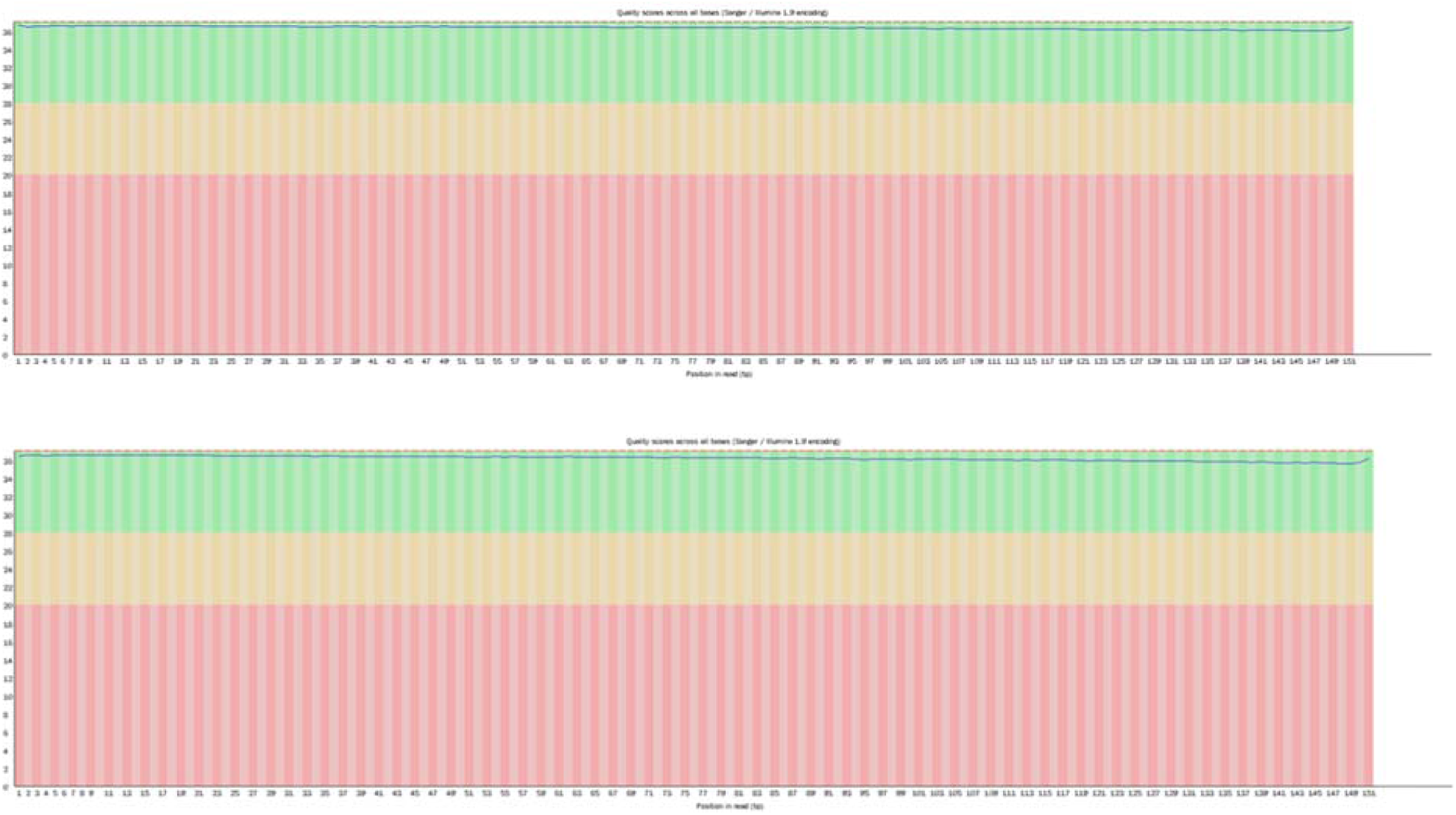
Figures of base quality of RNAtotal_1 (up: read; 1down: read2) at each cycle after filtering.

## References

Abrahamian, P., Cai, W., Nunziata, S.O., Ling, K-S., Jaiswal, N., Mavrodieva, V.A., et al. (2022). Comparative Analysis of Tomato Brown Rugose Fruit Virus Isolates Shows Limited Genetic Diversity. Viruses. 14(12): 2816. doi:10.3390/v14122816

Albrechtsen, S.E. (2006). Testing methods for seed transmitted viruses: principles and protocols. Publisher: CABI, Wallingford, 268 p.

Botermans, M., Pier, P. M., de Koning, Oplaat, C., Fowkes, A.R., McGreig, S., et al. (2023). Tomato Brown Rugose Fruit Virus Nextstrain Build Version 3: Rise of a Novel Clade. PhytoFrontiers. 2 (3): 442–446. doi: 10.1094/PHYTOFR-09-22-0090-A.

Çelik A, Coşkan S, Morca AF, Santosa AI, Koolivand D. (2022). Insight into population structure and evolutionary analysis of the emerging tomato brown rugose fruit virus. Plants 11:233279. doi: 10.3390/plants11233279.

Chung, B.N., Choi, K.S., Ahn, J.J., Joa, J.H., Do, K.S., Park, K.S. (2015). Effects of Temperature on Systemic Infection and Symptom Expression of Turnip mosaic virus in Chinese cabbage (*Brassica campestris*). Plant Pathol J. 31(4):363–70. doi: 10.5423/PPJ.NT.06.2015.0107.

Chellappan, P., Vanitharani, R., Ogbe, F., Fauquet, C.M. (2005). Effect of temperature on geminivirus-induced RNA Silencing in Plants. Plant Physiology. 138: 1828–1841. doi: 10.1104/pp.105.066563.

Cobos, A., Montes, N., López-Herranz, M., Gil-Valle, M., Pagán, I. (2019). Within-Host Multiplication and Speed of Colonization as Infection Traits Associated with Plant Virus Vertical Transmission. Journal of virology. 93(23): e01078–19. doi: 10.1128/JVI.01078-19.

Davino, S., Caruso, A.G., Bertacca, S., Barone, S., Panno, S. (2020). *Tomato Brown Rugose Fruit Virus*: Seed Transmission Rate and Efficacy of Different Seed Disinfection Treatments. Plants. 9: 1615. doi: 10.3390/plants9111615.

Dombrovsky, A., and Smith, E. (2017). Seed Transmission of Tobamoviruses: Aspects of Global Disease Distribution. InTech. doi: 10.5772/intechopen.70244

Dovas, C.I., Efthimiou, K., Katis, N.I. (2004). Generic detection and differentiation of tobamoviruses by a spot nested RT-PCR-RFLP using dI-containing primers along with homologous dG-containing primers. Journal of Virological Methods. 117: 137–144. doi: 10.1016/j.jviromet.2004.01.004.

Eichmeier, A., Hejlova, M., Orsagova, H., Frejlichova, L., Hakalova, E., Tomankova, K., et al. (2023). Characterization of Tomato Brown Rugose Fruit Virus (ToBRFV) Detected in Czech Republic. Diversity. 15(2):301. doi: 10.3390/d15020301.

Esmaeilzadeh, F., Santosa, A.I., Çelik, A., Koolivand, D. (2023). Revealing an Iranian Isolate of Tomato Brown Rugose Fruit Virus: Complete Genome Analysis and Mechanical Transmission. Microorganisms. 11(10):2434. doi: 10.3390/microorganisms11102434.

European and Mediterranean Plant Protection Organization (EPPO). (2019). Reporting Service 2019/192. https://gd.eppo.int/reporting/article-6622 (accessed 10 January 2022).

Fajinmi, A.A., and Fajinmi, O.B. (2010). Incidence of okra mosaic virus at different growth stages of okra plants (*Abelmoschus esculentus* (L.) Moench) under tropical conditions. Journal of General and Molecular Virology. 2: 028–031. http://www.academicjournals.org/JGMV

Fidan, H., Pelin, S., Kubra, Y., Bengi, T., Gozde, E., Ozer, C. (2020). Robust molecular detection of the new Tomato brown rugose fruit virus in infected tomato and pepper plant from Turkey. Journal of Integrative Agriculture. 19: 2–11. doi: 10.1016/S2095-3119(20)63335-4.

Froissart, R., Doumayrou, J., Vuillaume, F., Alizon, S., Michalakis, Y. (2010). The virulence – transmission trade-off in vector-borne plant viruses: a review of (non-)existing studies. Philosophical Transactions of the Royal Society B. 365: 1907–1918. doi: 10.1098/rstb.2010.0068.

García-Estrada, R.S., Diaz-Lara, A., Aguilar-Molina, V.H., Tovar-Pedraza, J.M. (2022). Viruses of Economic Impact on Tomato Crops in Mexico: From Diagnosis to Management-A Review. Viruses. 14(6):1251. doi: 10.3390/v14061251.

Hemmati, K., and McLean, D. L. (1977). Gamete-seed transmission of alfalfa mosaic virus and its effect on seed germination and yield in alfalfa plants. Phytopathology. 67: 576–579.

Herrera-Vásquez, J.A., Córdoba-Sellés, M.C., Cebrián, M.C., Alfaro-Fernandez, A., Jordá, C. (2009). Seed transmission of Melon necrotic spot virus and efficacy of seed-disinfection treatments. Plant Pathology. 58:7. doi: 10.1111/j.1365-3059.2008.01985.x.

Ishibashi, K. and Ishikawa, M. (2016). Replication of tobamovirus RNA. Annu. Rev. Phytopathol. 54:55–78. doi:10.1146/annurev-phyto-080615-100217.

Jordon-Thaden, I.E., Chanderbali, A.S., Gitzendanner, M.A., Soltis, D.E. (2015). Protocol note. Modified CTAB and Trizol protocols improve RNA extraction from chemically complex embryophyta. Applications in Plant Sciences. 3 (5): 1400105.

Lu, S., Wang, J., Chitsaz, F., Derbyshire, M.K., Geer, R.C., Gonzales, N.R., et al. (2020). CDD/SPARCLE: the conserved domain database in 2020, Nucleic Acids Research, 48: D265–D268. doi: 10.1093/nar/gkz991.

Luria, N., Smith, E., Reingold, V., Bekelman, I., Lapidot, M., Levin, I., et al. (2017). A New Israeli Tobamovirus Isolate Infects Tomato Plants Harboring Tm-22 Resistance Genes. PloS One. 12(1), e0170429. doi:10.1371/journal.pone.0170429

Magaña-Álvarez, A.A., Pérez-Brito, D., Vargas-Hernández, B.Y., Ramírez Pool, J.A., Núñez Muñoz, L.A., Salgado Ortiz, H., et al. (2021). Detection of *Tomato brown rugose fruit virus* (ToBRFV) in solanaceous plants in Mexico. J Plant Dis Prot. 128. 1627–1635. doi:10.1007/s41348-021-00496-1.

Marchler-Bauer, A., Bo, Y., Han, L., He, J., Lanczycki, C.J., Lu, S., et al. (2017). CDD/SPARCLE: functional classification of proteins via subfamily domain architectures. Nucleic Acids Research. 45: D200–D203. doi:10.1093/nar/gkw1129.

Marodin, J.C., Resende, F.V., Gabriel, A., Souza, R.J.De, Resende, J.T.V. De, Camargo, et al. (2019). Agronomic performance of both virus-infected and virus free garlic with different seed bulbs and clove sizes. Pesquisa Agropecuária Brasileira. 54: e01448. doi: 10.1590/S1678-3921.pab2019.v54.01448.

Matzrafi, M., Abu-Nassar, J., Klap, C., Shtarkman, M., Smith, E., Dombrovsky, A. (2023). *Solanum elaeagnifolium* and *S. rostratum* as potential hosts of the tomato brown rugose fruit virus. PLoS ONE 18(3): e0282441. doi: 10.1371/journal.pone.0282441

Mohan, B.G., Baruah, G., Sen, P., Deb, N.P., Kumar, B.B. (2020). “Chapter 11. Host-Parasite Interaction During Development of Major Seed-Transmitted Viral Diseases”. In Seed-Borne Diseases of Agricultural Crops: Detection, Diagnosis & Management, eds Kumar, R., Gupta, A. (Springer). 265–289. doi:10.1007/978-981-32-9046-4_11.

Ortiz-Martínez, L.E., Ochoa-Martínez, D.L., Rojas-Martínez, R.I., Aranda-Ocampo, S., Gómez Cruz, M.Á. (2022). Response of chili varieties to the infection with Tomato brown rugose fruit virus. Summa Phytopathologica. 4(4): 209–215. doi: 10.1590/0100-5405/250747.

Pagán I. (2022). Transmission through seeds: The unknown life of plant viruses. PLoS Pathog. 18(8):e1010707. doi: 10.1371/journal.ppat.1010707.

Panno S, Caruso, G.A., Stefano, B., Lo Bosco Giosuè, Ezequiel, R.A., Salvatore, D. (2020). Spread of *Tomato Brown Rugose Fruit Virus* in Sicily and Evaluation of the Spatiotemporal Dispersion in Experimental Conditions. Agronomy. 10: 834. doi: 10.3390/agronomy10060834.

Panno, S., Ruiz-Ruiz, S., Caruso, A.G., Alfaro-Fernandez, A., San Ambrosio, M.I., Davino, S. (2019). Real-time reverse transcription polymerase chain reaction development for rapid detection of *Tomato brown rugose fruit virus* and comparison with other techniques. PeerJ, 7: e7928. doi: 10.7717/peerj.7928.

Rizzo, D., Da, L.D., Panattoni, A., Salemi, C., Cappellini, G., Bartolini, L., et al. (2021). Rapid and Sensitive Detection of Tomato Brown Rugose Fruit Virus in Tomato and Pepper Seeds by Reverse Transcription Loop-Mediated Isothermal Amplification Assays (Real Time and Visual) and Comparison With RT-PCR End-Point and RT-qPCR Methods. Frontiers in Microbiology. 12:640932. doi: 10.3389/fmicb.2021.640932.

Rodríguez-Mendoza, J., García-Ávila, C.J., López-Buenfil, J.A., Araujo-Ruiz, K., Quezada-Salinas, A., Cambrón-Crisantos, J.M. (2019). Identification of Tomato brown rugose fruit virus by RT-PCR from a coding region of replicase (RdRP). Mexican Journal of Phytopathology. 37(2): 1–12. doi:10.18781/R.MEX.FIT.1902-6.

Sabra, A., Amer, M.A., Hussain, K., Zakri, A., Al-Shahwan, I.M., Al-Saleh, M.A. (2022). Occurrence and Distribution of Tomato Brown Rugose Fruit Virus Infecting Tomato Crop in Saudi Arabia. Plants (Basel). 11(22):3157. doi: 10.3390/plants11223157.

Salem, N., Mansour, A., Ciuffo, M., Falk, B.W., Turina, M. 2016. A new tobamovirus infecting tomato crops in Jordan. Arch. Virol. 161:2503–6. doi: 10.1007/s00705-015-2677-7.

Salem, N.M., Sulaiman, A., Samarah, N., Turina, M., Vallino, M. (2022). Localization and Mechanical Transmission of Tomato Brown Rugose Fruit Virus in Tomato Seeds. Plant Dis. 106(1):275–281. doi: 10.1094/PDIS-11-20-2413-RE.

Salem, N.M., Jewehan, A., Aranda M.A., Fox, A. (2023). Tomato Brown Rugose Fruit Virus Pandemic. Annual Review of Phytopathology. 61. doi:10.1146/annurev-phyto-021622-120703

Szittya, G., Silhavy, D., Molnár, A., Havelda, Z., Lovas, A., Lakatos, L., et al. (2003). Low temperature inhibits RNA silencing-mediated defense by the control of siRNA generation. EMBO J. 22: 633–640. doi: 10.1093/emboj/cdg74.

Vargas-Mejía, P., Rodríguez-Gómez, G., Salas-Aranda, D.A., et al. (2023). Identification and management of tomato brown rugose fruit virus in greenhouses in Mexico. Archives of Virology. 168(5):135. doi: 10.1007/s00705-023-05757-y.

Van de Vossenberg, B.T.L.H., Dawood, T., Woźny, M., Botermans, M. (2021). First Expansion of the Public Tomato Brown Rugose Fruit Virus (ToBRFV) Nextstrain Build; Inclusion of New Genomic and Epidemiological Data. PhytoFrontiers. 1(4): 359–363. doi: 10.1094/PHYTOFR-01-21-0005-A

Vasquez-Gutierrez, U., López, L.H.; Frías, T.G.A, Delgado, O.J.C., Flores, O.A., Aguirre, U.L.A. Hernández, J.A. (2024). Biological Exploration and Physicochemical Characteristics of Tomato Brown Rugose Fruit Virus in Several Host Crops. Agronomy. 14: 388. doi:10.3390/agronomy14020388.

Wang, J., Chitsaz, F., Derbyshire, M.K., Gonzales N.R., Gwadz, M., Shennan Lu, et al. (2023), The conserved domain database in 2023. Nucleic Acids Research. 51: D384–D388. doi:10.1093/nar/gkac1096

Zhang, X., Zhang, X., Singh, J., Li, D., Qua, F. (2012). Temperature-dependent survival of Turnip crinkle virus-infected Arabidopsis plants relies on an RNA silencing-based defense that requires DCL2, AGO2, and HEN1. J Virol. 12:6847–6854. doi: 10.1128/JVI.00497-12.

Zhi-yong, Y., Mei-sheng, Z., Hua-yu, M., Ling-zhi, L., Guang-ling, Y., Chao, G., Yanping, T., Xiang-dong, L. (2021). Biological and molecular characterization of tomato brown rugose fruit virus and development of quadruplex RT-PCR detection. Journal of Integrative Agriculture. 7 (20): 1871–1879. doi: 10.1016/S2095-3119(20)63275-0.

Zisi, Z., Ghijselings, L., Vogel, E., Vos, C., Matthijnssens, J. (2024). Single amino acid change in tomato brown rugose fruit virus breaks virus-specific resistance in new resistant tomato cultivar. Front Plant Sci. 7(15):1382862. doi: 10.3389/fpls.2024.1382862.

